# Testing the mettle of METAL: A comparison of phylogenomic methods using a challenging but well-resolved phylogeny

**DOI:** 10.1101/2024.02.28.582627

**Authors:** Edward L. Braun, Carl H. Oliveros, Noor D. White Carreiro, Min Zhao, Travis C. Glenn, Robb T. Brumfield, Michael J. Braun, Rebecca T. Kimball, Brant C. Faircloth

**Affiliations:** Department of Biology, University of Florida, Gainesville, Florida, USA, 32611; Department of Biological Sciences, Louisiana State University, Baton Rouge, LA, USA, 70803; Biological Imaging Core, National Eye Institute, National Institutes of Health, 9000 Rockville Pike, Bethesda, MD, USA, 20892; Department of Vertebrate Zoology, National Museum of Natural History, Smithsonian Institution, Washington, DC, USA, 20013; Department of Environmental Health Science and Institute of Bioinformatics, University of Georgia, 150 East Green St., Athens, GA, USA, 30602; Museum of Natural Science, Louisiana State University, Baton Rouge,LA, USA, 70803; Department of Biology and Biology Graduate Program, University of Maryland, College Park, MD 20742 USA

**Keywords:** Neighbor joining, Multispecies coalescent, Long branch attraction, Gene tree discordance

## Abstract

The evolutionary histories of different genomic regions typically differ from each other and from the underlying species phylogeny. This makes species tree estimation challenging. Here, we examine the performance of phylogenomic methods using a well-resolved phylogeny that nevertheless contains many difficult nodes, the species tree of living birds. We compared trees generated by maximum likelihood (ML) analysis of concatenated data, gene tree summary methods, and SVDquartets. We also conduct the first empirical test of a “new” method called METAL (Metric algorithm for Estimation of Trees based on Aggregation of Loci), which is based on evolutionary distances calculated using concatenated data. We conducted this test using a novel dataset comprising more than 4,000 ultraconserved element (UCE) loci from almost all bird families and two existing UCE and intron datasets sampled from almost all avian orders. We identified “reliable clades” very likely to be present in the true avian species tree and used them to assess method performance. ML analyses of concatenated data recovered almost all reliable clades with less data and greater robustness to missing data than other methods. METAL recovered many reliable clades, but only performed well with the largest datasets. Gene tree summary methods (weighted ASTRAL and weighted ASTRID) performed well; they required less data than METAL but more data than ML concatenation. SVDquartets exhibited the worst performance of the methods tested. In addition to the methodological insights, this study provides a novel estimate of avian phylogeny with almost 99% of the currently recognized avian families. Only one of the 181 reliable clades we examined was consistently resolved differently by ML concatenation versus other methods, suggesting that it may be possible to achieve consensus on the deep phylogeny of extant birds.

## Introduction

Advances in sequencing technologies and bioinformatics have revolutionized the field of phylogenetics, leading to the genesis of a novel field called phylogenomics (Philippe et al., 2005). Although phylogenomics originally referred to the use of phylogenetic methods to make functional inferences (Eisen, 1998), the term rapidly became associated with the use of large-scale sequence data matrices (often comprising hundreds or thousands of loci) for phylogenetic estimation. Three major findings emerged from early phylogenomic studies, two of which were evident in one of the earliest studies (Rokas et al., 2003). First, phylogenetic analyses using data from multiple genes often resulted in trees with very high support. Second, separate analyses of individual gene regions often produced conflicting trees. The third major finding, that trees generated using different analytical methods or data matrices can exhibit strongly-supported conflicts, emerged shortly thereafter (Jeffroy et al., 2006). Understanding the second and third findings has played a central role in the development of phylogenomics as a field.

The third discovery—the observation that phylogenomic studies can yield conflicting trees with high support—reflects the existence of systematic error (Delsuc et al., 2005; Jeffroy et al., 2006). The potential for systematic error in phylogenomics had long been clear based on theoretical studies (e.g., Felsenstein, 1978; Hendy and Penny, 1989) indicating that some phylogenetic methods can be inconsistent (i.e., they yield an incorrect estimate of phylogeny even with infinite data). Kim (2000) used tools from algebraic geometry to show why adding data to analyses that use inconsistent phylogenetic estimators results in increased support for an incorrect tree topology. Long-branch attraction is the best-characterized source of systematic bias (Felsenstein, 1978; Parks and Goldman, 2014; Susko and Roger, 2021), and it typically emerges when the model assumed by the analytical method is misspecified. More generally, the simplified models commonly used in phylogenomics are often unable to capture the full complexity of genome evolution, leading to the potential for bias (Gatesy, 2007). Although it is possible to implement more sophisticated models to avoid bias, the use of very complex models often pushes analyses of large datasets beyond the capabilities of available computational resources.

The second discovery, the existence of pervasive conflicts among gene trees, is potentially more complex and it contributes to the conflicts between estimates of species trees. Conflicts among individual gene trees arise in several ways. Stochastic error, where the data associated with individual genes are insufficient to yield an accurate estimate of their phylogeny (Chojnowski et al., 2008; Springer and Gatesy, 2016), is a simple explanation for apparent conflict. However, conflicts among gene trees may reflect biological sources of discordance. Maddison (1997) described the three causes of discordance among gene trees: incomplete lineage sorting (ILS), gene duplication followed by loss, and horizontal transfer. Of these, ILS is arguably the most important process because it is an unavoidable consequence of branching evolution given finite populations (Edwards, 2009; Rannala et al., 2020; Mirarab et al., 2021). The importance of the other two processes varies among lineages. Conflicts among gene trees can lead to systematic error in estimated species trees generated using the simplest analytical approach for phylogenomic data: maximum likelihood (ML) analysis of concatenated data matrices (Jeffroy et al., 2006; Kubatko and Degnan, 2007; Mirarab et al., 2021). The ML concatenation approach implicitly assumes a single underlying tree topology, which is typically a model misspecification because the data being concatenated reflect a mixture of gene trees that exhibit discordance (Edwards, 2009). This model misspecification can lead to ML concatenation converging on an incorrect estimate of the species tree in parts of parameter space (Roch and Steel, 2015).

Simulation studies that include ILS have shown that ML concatenation performs well in many parts of parameter space (Patel et al., 2013; Mirarab et al., 2016), although there are also parts of parameter space where it exhibits poor performance (Kubatko and Degnan, 2007). Finding that ML concatenation could be problematic led to development of multispecies coalescent (MSC) methods for species tree estimation that account for ILS in various ways. There are fully parametric MSC methods (Yang, 2002; Heled and Drummond, 2009), but the use of those methods in phylogenomics is limited by their computational requirements. Methods that accept a set of precomputed gene trees to produce an estimate of the species tree are much more practical in phylogenomic studies. ASTRAL, which generates an estimate of the species tree that agrees with the largest number of quartet trees induced by a set of input gene trees (Mirarab et al., 2014), is one of the most commonly used gene tree summary methods. An alternative way to summarize gene trees involves calculating the average distances between pairs of taxa in the gene trees and using that distance matrix to generate an estimate of the species tree; NJst (Liu and Yu, 2011) and ASTRID (Vachaspati and Warnow, 2015) are two implementations of this approach. Although gene tree summary methods are much faster than fully parametric methods, they can exhibit poor performance when the input gene trees have substantial estimation error (Patel et al., 2013; Gatesy and Springer, 2014; Meiklejohn et al., 2016; Springer and Gatesy, 2016; Zhao et al., 2023).

SVDquartets is an alternative MSC method that can use concatenated data matrices as input. Unlike ML concatenation, SVDquartets is a consistent estimator under fairly general conditions (Chifman and Kubatko, 2015; Wascher and Kubatko, 2021). It differs from gene tree summary methods in that it generates four-taxon trees based on the concatenated data and combines those quartets to generate the estimated species tree. The choice among the three possible trees for each quartet is based on the ranks of matrices populated with the 256 possible site pattern frequencies, where the ordering of the site pattern frequencies reflect the topology. The rank of the matrix for the appropriate quartet tree is *≤*10 in expectation. The use of simple matrix algebra makes SVDquartets computationally efficient for individual quartets while incorporating all sources of variability (i.e., both mutational variance and coalescent variance; see Huang et al., 2010) in the estimation process (Chifman and Kubatko, 2014). Because this method does not use estimated gene trees it cannot be affected by gene tree estimation error, a potential benefit for the method.

Dasarathy et al. (2015) proposed a different species tree estimator they called METAL; perhaps surprisingly, METAL is simply the neighbor-joining (NJ) method (Saitou and Nei, 1987) applied to a concatenated data matrix. The proof that METAL is a consistent estimator of the species tree (Dasarathy et al., 2015) assumed the data were generated under the Jukes and Cantor (1969) (JC) model, although they stated that METAL could be generalized for more complex models. Allman et al. (2019) extended the idea behind METAL to include NJ of logdet/paralinear distances (Lockhart et al., 1994; Lake, 1994); the logdet distance estimator is related to the general Markov model (Steel, 1994) so it is much more biologically-realistic than the JC model. In fact, Allman et al. (2019) showed that NJ of logdet distances was a consistent estimator of the species tree even when different parts of the genome evolved under different time-reversible substitution processes. Therefore, we suggest that the term METAL should be expanded to include NJ analyses of phylogenomic datasets more broadly. Indeed, the use of NJ is not critical; the proofs in Dasarathy et al. (2015) and Allman et al. (2019) show that the distances calculated using concatenated data matrices converge on the values expected given the true species tree. Thus, any distance method that yields the true tree given perfect distances (e.g., Fitch and Margoliash, 1967; Gascuel, 1997) will yield the true species tree in expectation (assuming the conditions described in the proofs are met). However, NJ has the benefit of being extremely computationally efficient, so it represents a desirable way to convert pairwise distances into trees.

Although NJ has been a popular method to generate trees using distances among taxa, it has fallen out of favor in the field of systematics. The use of NJ is more common when the tree topology is a nuisance parameter. For example, Poon et al. (2009) stated that NJ is a good way to generate trees for selection analyses because there is “…evidence that selection analyses are robust to some error in the phylogeny, hence a ‘quick and dirty’ method like NJ should be sufficient in most cases.” This perception of NJ seems at odds with the proofs (Liu and Edwards, 2009; Dasarathy et al., 2015; Allman et al., 2019) that it can be a consistent estimator of the species tree. However, consistency is an asymptotic property; it describes the expected behavior of an estimator in the limit of infinite data. Dasarathy et al. (2015) addressed the issue of data requirements, but it is difficult to draw firm conclusions for empirical studies from that theoretical work. Thus, it is important to evaluate the performance of METAL relative to other, commonly-used methods for phylogenomics in an empirical setting.

Here we examine the performance of multiple analytical methods, including METAL, using empirical phylogenomic datasets for birds. One criterion that can be used to evaluate the performance of analytical methods in empirical studies is the recovery of “reliable clades.” Obviously, it is impossible to know the true tree with certainty, making the definition of reliable clades subjective. Moreover, reliable clades that can be consistently defined may be too “easy” to recover and therefore uninformative with respect to performance because they are recovered by all methods. Even worse, it is possible for supposedly reliable clades to be falsified by other lines of evidence. For example, Li et al. (2001) assessed the performance of a new distance method using a “known” mammalian phylogeny that assumed an incorrect position for rodents, causing them to overstate the performance of their method (Braun, 2023). For these reasons, such tests should focus on lineages that have received intensive study, allowing identification of specific nodes that are challenging to recover but are nevertheless strongly corroborated by multiple lines of evidence. Relationships among the living families and orders of birds have received extensive study (Table 1). These studies have revealed many reliable clades at various depths in the avian tree; the recovery of those clades can be used to test the performance of phylogenetic methods.

**Table 1.** Analyses of deep avian phylogeny using multigene and phylogenomic datasets.

| Citation | # of Loci | Data Type(s) | # of Taxa<sup>2</sup> |
| --- | --- | --- | --- |
| **Multigene<sup>1</sup>** |  |  |  |
| Hackett et al. (2008) | 19 | Introns | 169 |
| Harshman et al. (2008) | 20 | Introns | 14 (Palaeognathae) |
| Haddrath and Baker (2012) | 27 | Coding exons and introns | 23 (Palaeognathae) |
| Wang et al. (2011) | 30 | Introns | 28 (Telluraves) |
| Kimball et al. (2013) | 31 | Introns | 77 |
| Smith et al. (2013) | 40 | Introns | 10 (Palaeognathae) |
| Reddy et al. (2017) | 54 | Introns | 235 |
| Liu et al. (2018) | 63 | Coding exons | 48 |
| **Phylogenomic<sup>1</sup>** |  |  |  |
| McCormack et al. (2013) | 1,541 | UCEs<sup>3</sup> | 32 (Neoaves) |
| Jarvis et al. (2014) | 11,839 | Whole genomes<sup>4</sup> | 48 |
| Baker et al. (2014) | 1,448 | Coding exons and UCEs | 198 |
| Prum et al. (2015) | 259 | Coding exons | 198 |
| Suh et al. (2015) | 2,118 | TE insertions<sup>5</sup> | 43 |
| Cloutier et al. (2019) | 20,850 | Whole genomes<sup>6</sup> | 15 (Palaeognathae) |
| White and Braun (2019) | 4,243 | UCEs | 23 (Strisores) |
| Kuhl et al. (2021) | —<sup>7</sup> | 3' UTRs<sup>8</sup> | 429 |
| Wang et al. (2022b) | —<sup>7</sup> | Whole genomes | 16 (Palaeognathae) |
| This study | 4,307 | UCEs | 394 |

<sup>1</sup> We define multigene studies as those with 11–100 loci and phylogenomic studies to those with >100 loci. Data type(s) reflect the majority of sequence data analyzed; many datasets include small amounts of other data types.
<sup>2</sup> This table is limited to studies that include multiple avian orders. Clade names in parentheses indicate that the taxon sample was limited to specific superordinal groups. Supraordinal clade names are defined below.
<sup>3</sup> Ultraconserved elements.
<sup>4</sup> Jarvis et al. \(2014\) reports separate and combined analyses of 8,253 coding exons, 2,516 introns (a subset of the 8,253 coding loci), and 3,586 UCEs along with an ML concatenation analyses of a 322 million bp whole genome alignment.
<sup>5</sup> Transposable element insertions.
<sup>6</sup> Cloutier et al. \(2019\) reports analyses of 12,676 conserved nonexonic elements (CNEEs; see Edwards et al., 2017, for a definition of this data type), 5,016 introns, 3,158 UCEs, and 4,274 TE insertions.
<sup>7</sup> Total number of loci sampled unspecified but clearly >100.
<sup>8</sup> 3' untranslated regions, largely obtained from transcriptome data.

Ideally, empirical tests of phylogenetic methods should use novel datasets that were not used to define the reliable target clades that will be used to evaluate method performance. For this study, we collected a novel dataset comprising 4,307 ultraconserved element (UCE) loci from 394 birds, representing all extant orders and almost all extant families of birds (as circumscribed by the IOC World Bird List v13.1; Gill et al., 2023). Although many phylogenomic studies of birds have used UCE data, most of those studies have focused on relationships within specific avian orders or families (e.g., Hosner et al., 2016; Manthey et al., 2016; Persons et al., 2016; Hosner et al., 2017; Oliveros et al., 2019a,b; Leite et al., 2021; Smith et al., 2022). Thus, efforts to define reliable clades have largely relied on non-UCE data (e.g., Hackett et al., 2008; Jarvis et al., 2014; Prum et al., 2015; Kuhl et al., 2021), making this novel UCE dataset largely independent of the studies used to define reliable clades. Consideration of the consistency and strength of support for clades in prior studies can be used to differentiate “easy” target clades (i.e., clades that any useful phylogenetic method will recover) from “hard” clades (i.e., clades that provide strong evidence that a method performs well). With the prior support for those clades in mind, we analyzed this novel UCE dataset using ML concatenation, gene tree summary methods (weighted ASTRAL and weighted ASTRID), SVDquartets, and METAL (using a variety of distance metrics). To further examine data requirements for METAL and the other methods, we also assessed their performance using the UCE and intron datasets from Jarvis et al. (2014), which had much larger numbers of aligned sites but much sparser taxon sampling (Table 1). These analyses allowed us to critically compare the relative performance of commonly used phylogenomic methods as well as the less commonly used METAL approach using a well-studied phylogeny where it is possible to assess the difficulty of recovering various nodes in the species tree.

## Methods

### DNA extraction, library preparation, and sequencing

We collected new UCE sequence data from 180 avian species and two crocodilian outgroups (*Alligator mississippiensis* and *Gavialis gangeticus*) using targeted enrichment (Faircloth et al., 2012). These taxa were chosen to mirror the sampling design of the “Early Bird” studies (Hackett et al., 2008; Reddy et al., 2017) and in many cases the samples we used corresponded to the same specimen or DNA sample. We acquired tissues for these taxa from vouchered specimens in major museum collections whenever possible (Table S1) and extracted DNA from various sources (blood, muscle tissue, liver tissue, and toepad clips from dried skins) using a phenol-chloroform technique (Rosel and Block, 1996) or DNeasy extraction kits (Qiagen, GmbH). We quantified DNA extracts using Qubit fluorometers (Life Technologies, Inc.). We sheared DNA extracted from muscle or liver to a length of approximately 300-600 bp using a Qsonica Q800R sonicator (Qsonica, L.L.C.) and 15–45 cycles of 20 seconds on and 20 seconds off at 25% amplitude. We did not shear DNA extracted from toepads.

We prepared sequencing libraries from sheared DNA in two ways. For the first method, we input 50-1,600 ng sheared DNA to a modified genomic DNA library preparation protocol (Kapa Biosystems, Inc.) that incorporated “with-bead” cleanup steps (Fisher et al., 2011) using a generic Solid Phase Reversible Immobilization (SPRI) substitute (Rohland and Reich, 2012). During ligation, we incorporated sample-specific single-indexed adapters to the ligation reaction (Faircloth and Glenn, 2012). Following ligation, we PCR-amplified the resulting libraries using HiFi HotStart polymerase (Kapa Biosystems, Inc.) and 10 PCR cycles following the manufacturer’s protocol. For the second method, we prepared libraries using Hyper Prep Kits (Kapa Biosystems, Inc.) following the manufacturer’s protocol at 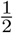 the recommended volume. We incorporated sample-specific dual-indexed adapters to library fragments (Glenn et al., 2019) by PCR amplifying the resulting libraries using HiFi Hotstart ReadyMix (Kapa Biosystems, Inc.) and 8-12 PCR cycles following the manufacturer’s protocol. After both ligation and amplification steps, we purified libraries prepared from toepad clips with a higher ratio (3x) of SPRI beads than recommended in the Hyper Prep Kit to retain small fragments.

We pooled samples in sets of eight at equimolar ratios prior to enrichment. Then we enriched UCE loci and performed post-enrichment amplification following standard protocols (Faircloth et al., 2012) using the Arbor Biosciences UCE 5Kv1 MyBaits Kit for tetrapods (Sun et al., 2014), which targets 5,060 UCE loci. This procedure involved hybridizing biotinylated RNA baits with pooled libraries for 24 h, capturing DNA bound to RNA baits using streptavidin-coated beads, and recovering enriched DNA with a 16-18 cycle PCR recovery step using HiFi Hotstart ReadyMix. After postenrichment amplification, we measured fragment size distributions using an Agilent 2100 Bioanalyzer and removed any remaining adapter-dimers using a 1.2x SPRI bead clean up. We quantified final libraries using a commercially available library quantification kit (Kapa Biosystems, Inc.), combined libraries into 10 nM pools, and sequenced the pooled libraries using either pairedend, 100 base pair (PE100) sequencing or paired-end, 150 base pair sequencing (PE150). The PE100 sequencing was conducted on a HiScan 1000 (George Washington University Medical School) or a HiSeq 2000 (UCLA Neuroscience Genomics Core) and the PE150 sequencing was conducted on a HiSeq 3000 (Oklahoma Medical Research Facility).

### Data quality control and assembly

We converted binary base call (BCL) files to FASTQ using the Illumina Casava ver. 1.8 software unless we received demultiplexed FASTQ data from the sequencing center. For all FASTQ files, we trimmed low-quality bases and adaptor sequences from reads using illumiprocessor ver. 2 (https://github.com/faircloth-lab/illumiprocessor) and assembled trimmed reads into contigs using Trinity ver. trinityrnaseq-r2013-02-25 (Grabherr et al., 2011).

We generated a 396 taxon dataset by combining the 182 new assemblies (including the two crocodilian outgroups) generated as described above with 214 assemblies from several different sources (Hosner et al., 2015; Manthey et al., 2016; Moyle et al., 2016; Campillo et al., 2018; Oliveros et al., 2019a,b). We used PHYLUCE (Faircloth, 2015) for the remaining data preparation steps following our standard protocols. Briefly, the remaining steps were the identification of contigs with UCE loci, collection of summary statistics, sequence alignment (which used MAFFT; Katoh and Standley, 2013), and alignment trimming, using Gblocks (Castresana, 2000) with the minimum number of sequences for a flank position (option “-b2”) set to 75%. After these steps, we assembled a data set with UCE loci that were present in at least 75% of all 396 taxa (see Table S3 for details regarding the data recovery). This taxon sample includes representatives of all 44 orders and 250 families of extant birds recognized by the IOC World Bird List (Table S2); only three families recognized by that taxonomy are absent. All three missing families are small; Pluvianellidae is monotypic, Alcippeidae includes one genus with ten species, and Teretistridae includes one genus with two species. The dataset comprised 4,307 loci and the mean alignment length for each locus was 474 bp (range 120–1,465 bp with a median length of 470 bp). The maximum amount of missing data and gaps for any taxon was extremely high (Table 2), so we constructed a second data matrix where the number of resolved bases in all taxa was >75% of the number of bases in the taxon with maximum number of resolved bases. This only resulted in the deletion of 27 taxa (Table S3), all of which also had short contigs and, in most cases, fewer recovered loci. This resulted in two datasets, a complete “allfam” dataset with 396 taxa and a “well-sampled allfam” dataset with 369 taxa (Table 2).

**Table 2.** Dataset sizes and characteristics.

| Dataset<sup>1</sup> | Aligned Sites | Parsimony<br>Informative sites | Mean | Gaps and Missing Data<sup>2</sup> |  |  |
| --- | --- | --- | --- | --- | --- | --- |
|  |  |  |  | Minimum | Median | Maximum |
| allfam (all) | 2,042,823 | 852,768 | 9.33% | 1.46% | 23.32% | 81.15% |
| allfam (ws) | 2,042,823 | 830,321 | 6.43% | 1.46% | 4.98% | 25.26% |
| Jarvis UCE | 9,229,346 | 3,848,788 | 19.11% | 16.79% | 18.26% | 33.43% |
| Jarvis intron | 19,258,311 | 10,777,039 | 30.02% | 20.60% | 29.25% | 45.72% |

<sup>1</sup> “allfam (all)” refers to the complete allfam dataset (396 taxa) and “allfam (ws)” refers to the well-sampled subset (369 taxa) of the allfam dataset.
<sup>2</sup> Percentage of gaps and missing data per taxon. This includes sites masked by TAPER.

To explore the importance of dataset size, we also reanalyzed the two non-coding datasets from Jarvis et al. (2014) that had more sequence data but fewer taxa. These datasets comprise UCEs and introns (hereafter called the “Jarvis UCE” and “Jarvis intron” datasets) from 48 taxa that represent 35 (of 44) living bird orders based on the IOC World Bird List. The Jarvis UCE dataset comprises 3,679 loci and the mean length of individual alignments is *∼*2.5 kbp. The Jarvis intron dataset comprised introns from 2,516 genic loci. The individual alignments typically include multiple introns from each transcription unit, leading to a mean alignment length that was much larger than the other datasets (*∼*7.7 kbp) and a very skewed distribution of intron alignment lengths (the range was 80 bp to *∼*74.7 kbp and the median alignment length was *∼*4.9 kbp). We downloaded the trimmed alignments (see Jarvis et al., 2015, for details regarding data availability) and masked potentially problematic sequences using TAPER (Zhang et al., 2021). TAPER masking was conducted because Springer and Gatesy (2017) identified some non-homologous regions in the alignments; Zhang et al. (2021) showed that TAPER identified at least part of the erroneous sequences in almost half of the 68 cases highlighted by Springer and Gatesy (2017) and most of the remaining homology errors were short. TAPER masks individual sequences so it is not overly aggressive in sequence filtering. This approach allowed us to test all methods with “cleaner” datasets without reducing the overall number of sites. Concatenated files for the Jarvis datasets were generated using simple_concat.pl, available from https://github.com/ebraun68/RYcode.

We converted binary base call (BCL) files to FASTQ using the Illumina Casava ver. 1.8 software unless we received demultiplexed FASTQ data from the sequencing center. For all FASTQ files, we trimmed low-quality bases and adaptor sequences from reads using illumiprocessor ver. 2 (https://github.com/faircloth-lab/illumiprocessor) and assembled trimmed reads into contigs using Trinity ver. trinityrnaseq-r2013-02-25 (Grabherr et al., 2011).

We generated a 396 taxon dataset by combining the 182 new assemblies (including the two crocodilian outgroups) generated as described above with 214 assemblies from several different sources (Hosner et al., 2015; Manthey et al., 2016; Moyle et al., 2016; Campillo et al., 2018; Oliveros et al., 2019a,b). We used PHYLUCE (Faircloth, 2015) for the remaining data preparation steps following our standard protocols. Briefly, the remaining steps were the identification of contigs with UCE loci, collection of summary statistics, sequence alignment (which used MAFFT; Katoh and Standley, 2013), and alignment trimming, using Gblocks (Castresana, 2000) with the minimum number of sequences for a flank position (option “-b2”) set to 75%. After these steps, we assembled a data set with UCE loci that were present in at least 75% of all 396 taxa (see Table S3 for details regarding the data recovery). This taxon sample includes representatives of all 44 orders and 250 families of extant birds recognized by the IOC World Bird List (Table S2); only three families recognized by that taxonomy are absent. All three missing families are small; Pluvianellidae is monotypic, Alcippeidae includes one genus with ten species, and Teretistridae includes one genus with two species. The dataset comprised 4,307 loci and the mean alignment length for each locus was 474 bp (range 120–1,465 bp with a median length of 470 bp). The maximum amount of missing data and gaps for any taxon was extremely high (Table 2), so we constructed a second data matrix where the number of resolved bases in all taxa was *>*75% of the number of bases in the taxon with maximum number of resolved bases. This only resulted in the deletion of 27 taxa (Table S3), all of which also had short contigs and, in most cases, fewer recovered loci. This resulted in two datasets, a complete “allfam” dataset with 396 taxa and a “well-sampled allfam” dataset with 369 taxa (Table 2).

To explore the importance of dataset size, we also reanalyzed the two non-coding datasets from Jarvis et al. (2014) that had more sequence data but fewer taxa. These datasets comprise UCEs and introns (hereafter called the “Jarvis UCE” and “Jarvis intron” datasets) from 48 taxa that represent 35 (of 44) living bird orders based on the IOC World Bird List. The Jarvis UCE dataset comprises 3,679 loci and the mean length of individual alignments is *∼*2.5 kbp. The Jarvis intron dataset comprised introns from 2,516 genic loci. The individual alignments typically include multiple introns from each transcription unit, leading to a mean alignment length that was much larger than the other datasets (*∼*7.7 kbp) and a very skewed distribution of intron alignment lengths (the range was 80 bp to *∼*74.7 kbp and the median alignment length was *∼*4.9 kbp). We downloaded the trimmed alignments (see Jarvis et al., 2015, for details regarding data availability) and masked potentially problematic sequences using TAPER (Zhang et al., 2021). TAPER masking was conducted because Springer and Gatesy (2017) identified some non-homologous regions in the alignments; Zhang et al. (2021) showed that TAPER identified at least part of the erroneous sequences in almost half of the 68 cases highlighted by Springer and Gatesy (2017) and most of the remaining homology errors were short. TAPER masks individual sequences so it is not overly aggressive in sequence filtering. This approach allowed us to test all methods with “cleaner” datasets without reducing the overall number of sites. Concatenated files for the Jarvis datasets were generated using simple_concat.pl, available from https://github.com/ebraun68/RYcode.

### Phylogenetic analyses

We conduced ML concatenation analyses using two different programs: IQ-TREE v. 2.2.2.7 (Minh et al., 2020) and ExaML ver. 3.0.15 (Kozlov et al., 2015). In IQTREE we identified the best-fitting model using the -m TEST option (Kalyaanamoorthy et al., 2017), which examines a set of models comprising the general time reversible model with invariant sites and Γ-distributed rates (the GTR+I+Γ model) and its submodels. We used the ultrafast bootstrap (Hoang et al., 2018) with 1000 replicates to assess support in the IQ-TREE analyses. We conducted the ExaML analysis to determine whether our ML concatenation analyses were sensitive to the program used for the ML searches; we assumed the GTR+Γ model in the ExaML analysis and we evaluated node support using 100 bootstrap replicates. We tested for convergence of bootstrap replicates using a posteriori “bootstopping” (Pattengale et al., 2009) (Pattengale et al. 2009) using the ‘autoMRE’ option in RAxML 8.2.8 (Stamatakis, 2014). We mapped bootstrap support onto the ExaML tree using sumtrees.py from the DendroPy package (Sukumaran and Holder, 2010). We found that the IQ-TREE and ExaML trees for the complete allfam dataset had very similar topologies, so we only used IQ-TREE for analyses of the other datasets because it was the faster program.

We also used IQ-TREE to generate ML estimates of gene trees for summary coalescent analyses; as above, we used the -m TEST option to identify the best-fitting model and we used the parametric approximate likelihood ratio test (the –alrt 0 option) (Anisimova and Gascuel, 2006) to assess branch support in the gene trees. The gene trees with support values were used as input for the weighted versions of ASTRAL (wASTRAL, Zhang and Mirarab, 2022) and ASTRID (wASTRID, Liu and Warnow, 2023). We assessed support in wASTRAL using the local posterior probabilities (Sayyari and Mirarab, 2016) generated by that program. wASTRID does not produce support values, so we generated 100 sets of input gene trees by sampling gene trees with replacement using wastral_boot.pl, available from https://github.com/ebraun68/METALtest. Note that all of the gene trees had –alrt 0 support values that are used by wASTRID and those support values were not changed as part of this bootstrapping. We used sumtrees.py to map bootstrap support onto the wASTRID trees.

We used PAUP* ver. 4.0a build 166 (Swofford, 2003) for SVDquartets analyses, sampling 10^7^ random quartets for the allfam datasets and all quartets for the Jarvis UCE and intron datasets. We also conducted an SVDquartets analysis of the well-sampled allfam dataset with exhaustive quartet sampling by dividing the taxa into two subsets (passerines and non-passerines, the former with a small number of non-passerine outgroups and the latter with a small number of passerines to allow us to combine the two subtrees). The complete quartet sampling subtrees were combined to generate a complete SVDquartets tree using the MRP method (Baum, 1992; Ragan, 1992). We used clann (Creevey and McInerney, 2005) to code the trees (including a “taxonomic backbone” tree) for MRP analysis and PAUP* to identify the most parsimonious tree that combined the two trees. We used the bootstrap with 100 replicates to assess support for all SVDquartets analyses that used complete quartet sampling. We used bootstrapping to assess support for the SVDquartets trees, mapping values onto the SVDquartets trees using sumtrees.py.

We used PAUP* for all METAL analyses; almost all of the analyses used NJ to generate the trees (the exception was a test of the BioNJ method described by Gascuel, 1997). We tested a standardized set of distance metrics with all four datasets: 1) *p*-distances; 2) ML distances; 3) “trace” distances; and 4) logdet distances (see Appendix 1 for descriptions of the distance estimators). Trace distances and ML distances used the best-fitting model (and among-sites rate heterogeneity parameters) identified by ML in IQ-TREE (using the -m TEST option). ML distances used estimates of the rate matrix, state frequency, and rate heterogeneity parameters from IQ-TREE. Trace distances only use rate heterogeneity parameters; they were also set at the values of the IQ-TREE estimates. Logdet distances cannot be combined with Γ-distributed rates across sites, so we tested two different site deletion strategies. We call logdet distances based on all sites “logdet i.r.” (identical rates); this distance assumes sites have the same underlying rates (i.e., there is no among-sites rate heterogeneity). We call logdet with invariant site deletion “logdet-inv” distances; we used the ML estimate of the invariant site proportion from IQ-TREE (when the best-model included +I) for this calculation. Finally, we call logdet distances based on parsimony informative site “logdet inf sites” distances; to calculate these distances we excluded the parsimony uninformative sites in PAUP* before conducting the distance calculations. We assessed support in METAL analyses using the bootstrap with 100 replicates; bootstrap support was mapped onto the NJ trees using sumtrees.py, as described above.

To better examine the dataset size requirement for METAL (while also controlling for data type) we randomly subsampled the Jarvis intron dataset (which is the largest dataset) to generate data subsets with numbers of parsimony informative sites similar to the Jarvis UCE dataset. We used the program shuffle_locuslist.pl (available from https://github.com/ebraun68/compdisttest) to randomly sample 900 intron alignments without replacement to generate each subset as described in Braun (2023). Since complete intron alignments were sampled, this resulted in datasets of different sizes; we provide the precise dataset sizes when we report results for each dataset. We estimated that 900 intronic alignments would yield datasets with numbers of informative sites comparable to the Jarvis UCE dataset. NJ analyses of the random subsamples were limited to trace distances (with the GTR+Γ model) and logdet inf sites distances.

We used the AssessMonophyly function in R package MonoPhy (Schwery and O’Meara, 2016) to assess monophyly of superorders, orders, superfamilies and families and summarize the number of those clades present in each tree. The R package ape (Paradis and Schliep, 2019) was used to process the trees.

### Identification of reliable clades

The most fundamental property of any “good” phylogenetic method is the ability, when given sufficient data, to yield an estimate of phylogeny that includes most or all of the clades present in the true species tree. Although parts of the bird tree remain highly uncertain (Suh, 2016; Braun et al., 2019), many avian clades are strongly corroborated by multiple lines of genetic or genomic evidence (such as the studies in Table 1). In some cases, analyses of morphological data support these same clades (e.g., Livezey and Zusi, 2007; Mayr, 2010). These strongly corroborated clades can be considered ‘reliable’ and should be recovered by any analytical method well-suited to the task. Thus, we use reliable clade recovery—the number of those reliable clades present in trees generated using different analytical methods—as a metric to evaluate performance.

We divided the reliable clades into 12 categories based on two characteristics: their relative ease of recovery and their depth in the tree (Table 3). More specifically, we used three “ease” of clade recovery categories and four clade depth categories. The ease of recovery categories are: 1) “easy” clades are recovered in a large proportion (at least 50%) of individual gene trees; 2) “medium” clades require multigene datasets for recovery; and 3) “hard” clades that require phylogenomic datasets for recovery. As in Table 1, we viewed multigene datasets as those with 11–100 loci and phylogenomic datasets as those with *>*100 loci. We expect any useful analytical method to yield a tree that includes the easy clades, whereas the hard clades may only emerge when analytical methods exhibit very good performance.

**Table 3.** Relative ease of recovery for reliable clades.

|  | Easy | Medium | Hard | TOTAL |
| --- | --- | --- | --- | --- |
| Families<sup>1</sup> | 93 | — | — | 93 |
| Superfamilies<sup>2</sup> | 3 / 1 | 14 | 4 | 21 / 1 |
| Orders<sup>1,2</sup> | 36 / 1 | 3 | — | 39 / 1 |
| Superorders<sup>2</sup> | 7 / 5 | 12 / 11 | 9 / 8 | 28 / 24 |
| TOTAL<sup>2</sup> | 139 / 7 | 29 / 11 | 13 / 8 | 181 / 26 |

<sup>1</sup> The complete allfam dataset includes taxa from 250 families and 44 orders, but we only consider families with at least two taxa sampled in this table. Ten of the 93 families sampled for two or more taxa in the complete allfam dataset were reduced to a single representative in the well-sampled allfam dataset.
<sup>2</sup> Numbers after the slashes correspond to the reliable clades we scored for analyses of the Jarvis datasets, which had limited taxon sampling (48 taxa). The only superfamily we scored in the Jarvis trees was Eupasseres and the only order we scored was Passeriformes.

To define reliable clades as objectively as possible we restricted our consideration to clades that 1) have published names with clear circumscriptions and 2) are supported by the lines of evidence described below. For families, we used the circumscriptions from IOC World Bird List v. 13.1 (Gill et al., 2023). The IOC World Bird List is one of four taxonomies used by the ornithological community (see Hosner et al., 2022, for additional details); we chose the IOC World Bird List because it is updated regularly and the ordinal circumscriptions are completely consistent with recent phylogenomic studies. The IOC familial circumscriptions have received extensive evaluation by experts, but the evidence supporting them often comes from studies that use small numbers of nuclear loci. Therefore, we classified all but one of the 94 families that we sampled for at least two taxa in the allfam dataset as easy clades. The exception is the IOC circumscription of Tityridae (tityras, mourners, and allies), which we do not view as reliable. Indeed, some taxonomic treatments already split Tityridae into multiple families; see, for example, the South American Classification Committee proposal 827 and note that Oliveros et al. (2019b) also treated Tityridae as non-monophyletic.

The identification of reliable suprafamilial clades was more challenging because specific authorities do not exist for this taxonomic level. We chose to focus on circumscriptions of superfamilies in Oliveros et al. (2019b), which were limited to the largest avian order, Passeriformes, where we sampled 143 of the 145 IOC families. We evaluated the reliability and ease of recovery for the 22 suprafamilial clades labeled in the Oliveros et al. (2019b) tree by determining whether the 22 clades were present in the Burleigh et al. (2015) and Kuhl et al. (2021) trees. Burleigh et al. (2015) is a supermatrix tree of 22 nuclear loci and seven mitochondrial regions); thus, we view it as comparable to a multigene study. Kuhl et al. (2021) is a phylogenomic study (Table 1). Of the 22 suprafamilial clades we considered, 17 were present in both the Burleigh et al. (2015) and Kuhl et al. (2021) trees. We placed three of those suprafamilial clades (Eupasseres, Tyranni, and Passeri) in the easy category based on earlier studies that used individual gene regions Ericson et al. (2002); Barker et al. (2004). We placed three suprafamilial clades in the hard category because they are present in the Kuhl et al. (2021) tree but not the Burleigh et al. (2015) tree and excluded one (Locustelloidea as circumscribed in Oliveros et al., 2019b) because it was not recovered in the Burleigh et al. (2015) tree or the Kuhl et al. (2021) tree. We placed the remaining suprafamilial clades in the medium category.

For orders we used the IOC World Bird List circumscriptions Gill et al. (2023). All of the currently recognized IOC orders have received support in multigene studies like Hackett et al. (2008) and Reddy et al. (2017); the question is whether they represent easy or medium clades. To examine that question we inspected the 56 individual gene trees from Reddy et al. (2017), finding that 38 IOC orders were present in *≥*50% of those gene trees and three were present in *<*50% of those gene trees (Table S4). The 38 orders supported in most individual gene trees were classified as easy clades and the three that were not (Pelecaniformes, Accipitriformes, and Piciformes) were classified as medium clades. We could not assess monophyly for three IOC orders because they are monotypic (Leptosomiformes and Opisthocomiformes) or sampled for a single species in the allfam dataset (Struthioniformes).

Our evaluation of supraordinal clades was guided by the work of Sangster et al. (2022). The primary goal of that study was to propose formal names for supraordinal bird clades and register those names with the PhyloCode (Cantino and de Queiroz, 2020), an authority for phylogenetic taxonomy. Sangster et al. (2022) wanted to limit the registered names to clades that are very likely to be present in the true avian species tree, so they evaluated supraordinal clades in a standardized manner. Specifically, they chose a standardized set of ten studies for Neoaves and a second standardized set of 12 studies for Palaeognathae and registered names for clades supported by at least four of those studies, focusing on the trees preferred by the authors of each study. In addition to their full consideration of the standardized sets of studies, Sangster et al. (2022) also consulted other publications that did not present a preferred phylogeny and/or focused on reanalyses of already published data. In this study, we break the named supraordinal clades into the three difficulty categories. Easy supraordinal clades meet two criteria: 1) they are supported by all studies evaluated by Sangster et al. (2022) *and* 2) they are found in *≥*50% of trees based on individual gene regions using the results in Fig. 1b of Hackett et al. (2008) and Fig. 5c of Jarvis et al. (2014) to establish whether clades met the 50% threshold for individual gene tree support (we focused on non-coding gene trees in Jarvis et al., 2014, because the trees based on coding data are poorly supported). Medium supraordinal clades were supported by all studies evaluated Sangster et al. (2022) and hard supraordinal clades were supported by at least four of the ten studies evaluated by Sangster et al. (2022). A consensus phylogeny of orders and superorders and their difficulty categories is presented in Figure 1. Agreement between trees based on nucleotide data and transposable element (TE) insertions provide especially strong corroboration for clades because the processes by which nucleotide sequences and TE insertions evolve are very different, as are the assumptions made by phylogenetic analyses using those data types. Therefore, we indicate the names of clades with support from at least five TE insertions in Suh et al. (2015) or Simmons et al. (2022) using bold text in all figures (e.g., Fig. 1b).

**Figure 1.**
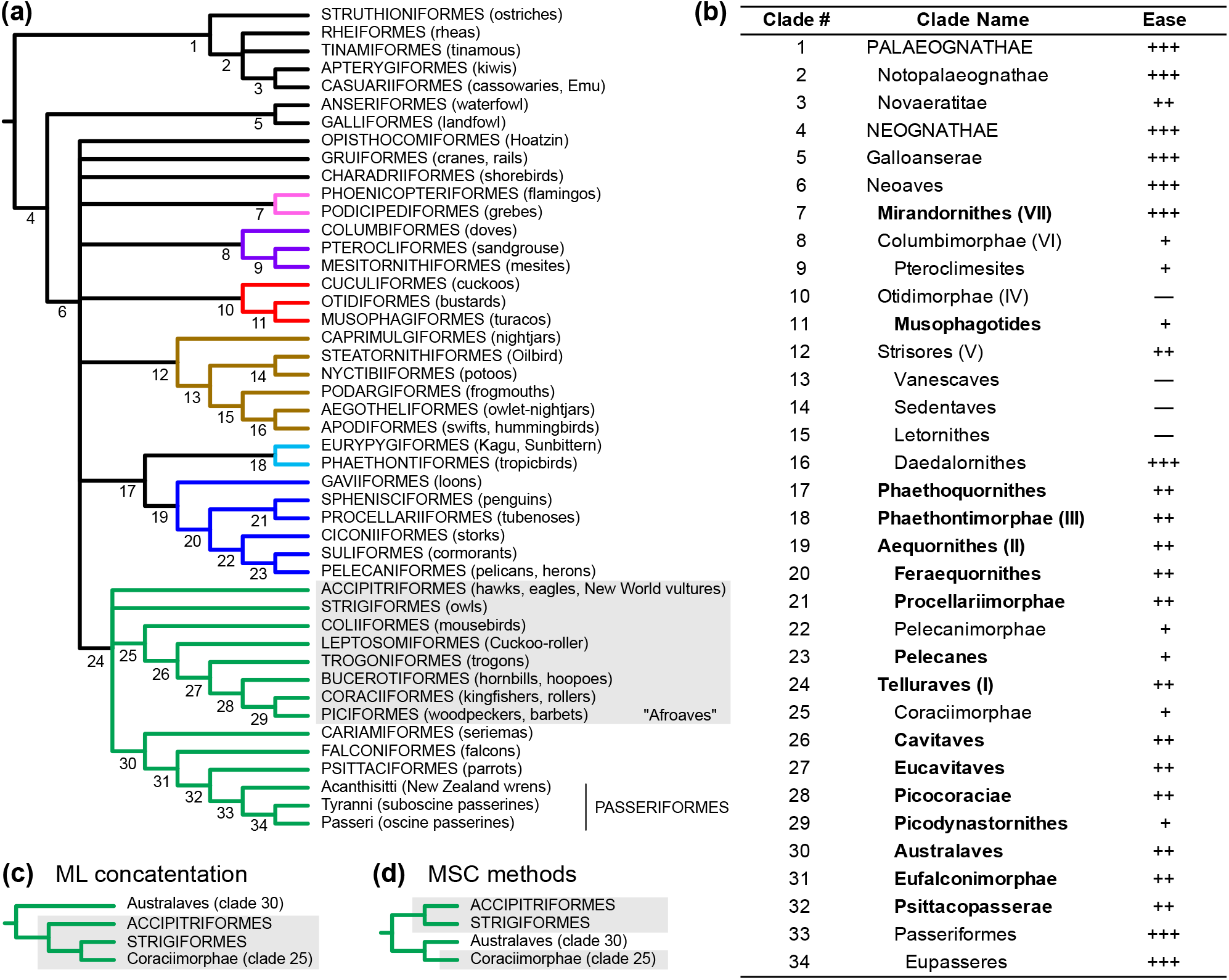
Relationships among avian orders based on published studies. (**a**) Cladogram based on Sangster et al. (2022) with minor modifications (see text). Numbers below nodes correspond to the clade names in (**b**) and colors indicate the “magnificent seven” clades defined by Reddy et al. (2017). Shaded taxa correspond to “Afroaves” *sensu* Ericson (2012). (**b**) Ease of recovery for reliable supraordinal clades in (**a**). ‘+++’ for easy clades, ‘++’ for medium clades, or ‘+’ for hard clades. Clades with a ‘—’ are either unreliable or their reliability was not assessed. Clades with bold names had support from at least five TE insertions in Suh et al. (2015) or Simmons et al. (2022). Roman numerals after the names of clades 7, 8, 10, 12, 18, 19, and 24 reflect the numbering of the magnificent seven clades in Reddy et al. (2017) (**c**) Telluraves topology typically recovered in ML concatenation analyses; Afroaves is shaded. (**d**) Telluraves topology recovered in many MSC analyses, examples include Fig. 3b in Jarvis et al. (2014) and Fig. S16 in Zhang and Mirarab (2022); Afroaves is non-monophyletic given the “MSC topology.”

Relationships among a few taxa that have been resolved one way in ML concatenation analyses and a different way by MSC methods (typically gene tree summary methods) in earlier analyses are of particular interest. These clades should not be included in our set of reliable clades because published information suggests that their recovery is method dependent, so we treat them as uncertain. The relevant clades are three supraordinal clades (some of which are mutually exclusive) and one suprafamilial clade. At the supraordinal level, ML concatenation typically supports Afroaves (Fig. 1c) whereas MSC methods often support placement of an Accipitriformes+Strigiformes clade sister to Coraciimorphae+Australaves (Fig. 1d) (note that Coraciimorphae+Australaves contradicts Afroaves). Within passerines, Oliveros et al. (2019b) found that the position of Calyptophilidae (a family comprising two species, herein represented by *Calyptophilus tertius*, the Western Chat-Tanager) exhibited a consistent pattern of conflict (i.e., all MSC trees agree and differ from the ML concatenation tree). Recovery of these clades has the potential to provide information about method performance because they are candidates for cases where ML concatenation might be biased due to the MSC.

### *A posteriori* tests of reliable clades

After novel phylogenomic tree with 124 birds sampled from almost all of the orders recognized in the IOC taxonomy (Wu et al., 2024) was published after we defined these reliable clades. This novel publication allowed us to conduct another test the of hypothesis that the reliable clades we defined are present in the true avian species tree. The Wu et al. (2024) study used a total of 25,460 loci extracted from whole genomes (5,756 coding regions, 12,449 CNEEs, 4,871 introns, and 2,384 intergenic segments). The taxon sampling of Wu et al. (2024) clades allowed us to evaluate the the reliable supraordinal clades and three passerine superfamiles (Eupasseres, Tyranni, and Passeri); the number of taxa sampled within each order was not sufficient to provide a rigorous test of monophyly of the orders. Wu et al. (2024) present an NJst (Liu and Yu, 2011) tree based on gene trees for a subset of their (the coding exons, introns, and intergenic segments after removal of topological outlier gene trees) in the main paper as their “primary tree”; they also present two trees for each data type in their supporting material (an NJst tree and an ML concatenation tree). We assessed the presence or absence of our reliable clades.

With the exception of Novaeratitae, all reliable clades in Fig. 1b were present in the primary Wu et al. (2024) tree (Supplementary Fig. S1). Most of the uncertain clades were also present; specifically, Accipitriformes+Strigiformes, Otidimorphae (clade 10), and the three uncertain nodes within Strisores (Vanescaves, Sedentaves, and Letornithes, clades 13–15) were all present. However, the Accipitriformes+Strigiformes clade was not sister to the remaining Telluraves (i.e., they did not recover Coraciimorphae+Australaves; see Fig. 1d). Instead, Wu et al. (2024) recovered a clade comprising Australaves, Accipitriformes, and Strigiformes, which they named Hieraves. The recovery of reliable clades was more variable in the separate analyses of individual data types (Supplementary Fig. S1), although we note that the number of clades recovered in each of our categories (easy, medium, and hard) was consistent with our classification. The analyses of introns and CNEEs recovered the largest numbers of reliable clades. The observation that analyses of introns and CNEEs recovered more reliable clades than analyses of coding sequences is consistent with the results of Reddy et al. (2017) and Braun and Kimball (2021).

## Results

### ML concatenation is more robust to missing data than MSC methods

Concatenated ML analyses of the complete allfam UCE dataset recovered the vast majority of reliable clades, yielding trees (e.g., Fig. 2 and File S1) that are quite similar to the trees from earlier phylogenomic studies (e.g., Jarvis et al., 2014; Kuhl et al., 2021). Both IQTREE and ExaML recovered 28 of 30 reliable supraordinal clades (Fig 3a). All orders, including the three medium difficulty orders, and all reliable superfamilies were monophyletic (Fig. 3b,c), as were all families designated as reliable. Thus, of 181 reliable clades addressed by the allfam dataset, concatenated ML recovered 179 of them, including all easy, all but one medium, and all but one of the hard clades.

**Figure 2.**
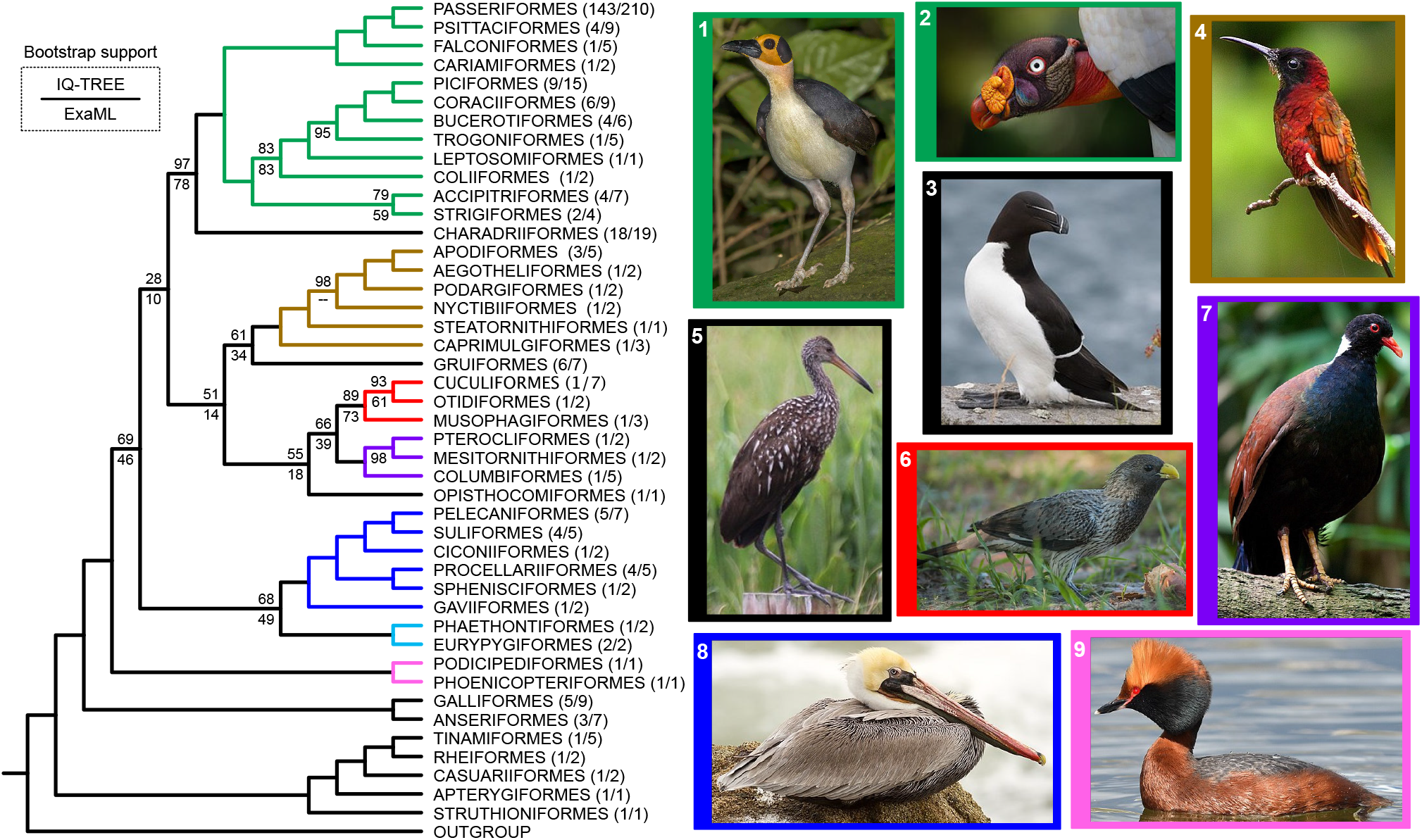
ML concatenation tree for the complete allfam dataset. Relationships among orders based on IQ-TREE are shown. IQ-TREE ultrafast bootstrap support values are presented above branches and ExaML bootstrap support values are below branches. Unannotated branches had complete (100%) support. There was only one conflict between the ML trees generated using the two programs at the level of orders; Steatornithiformes+Nyctibiiformes (clade 14, Sedentaves) is present in the ExaML tree (with 62% bootstrap support) but absent in this tree (the ExaML tree is available in File S1). The numbers of families and species sampled from each order are presented in parentheses after each order. Images to the right of the tree illustrate the diverse phenotypes of birds, with the frames for each image corresponding to the clade colors in the tree. The species in the images are: 1) *Picathartes gymnocephalus* (White-necked Rockfowl, Passeriformes: Picathartidae); 2) *Sarcoramphus papa* (King Vulture, Accipitriformes: Cathartidae); 3) *Alca torda* (Razorbill, Charadriiformes: Alcidae); 4) *Topaza pella* (Crimson Topaz, Apodiformes: Trochilidae); 5) *Aramus guarauna* (Limpkin, Gruiformes: Aramidae) ; 6) *Crinifer piscator* (Western Plantain-eater, Musophagiformes: Musophagidae); 7) *Otidiphaps nobilis* (Pheasant Pigeon, Columbiformes: Columbidae); 8) *Pelecanus occidentalis* (Brown Pelican, Pelecaniformes: Pelecanidae); and 9) *Podiceps auritus* (Horned Grebe, Podicipediformes: Podicipedidae). Images are from Wikimedia Commons, their sources and license information are available in Appendix 2.

**Figure 3.**
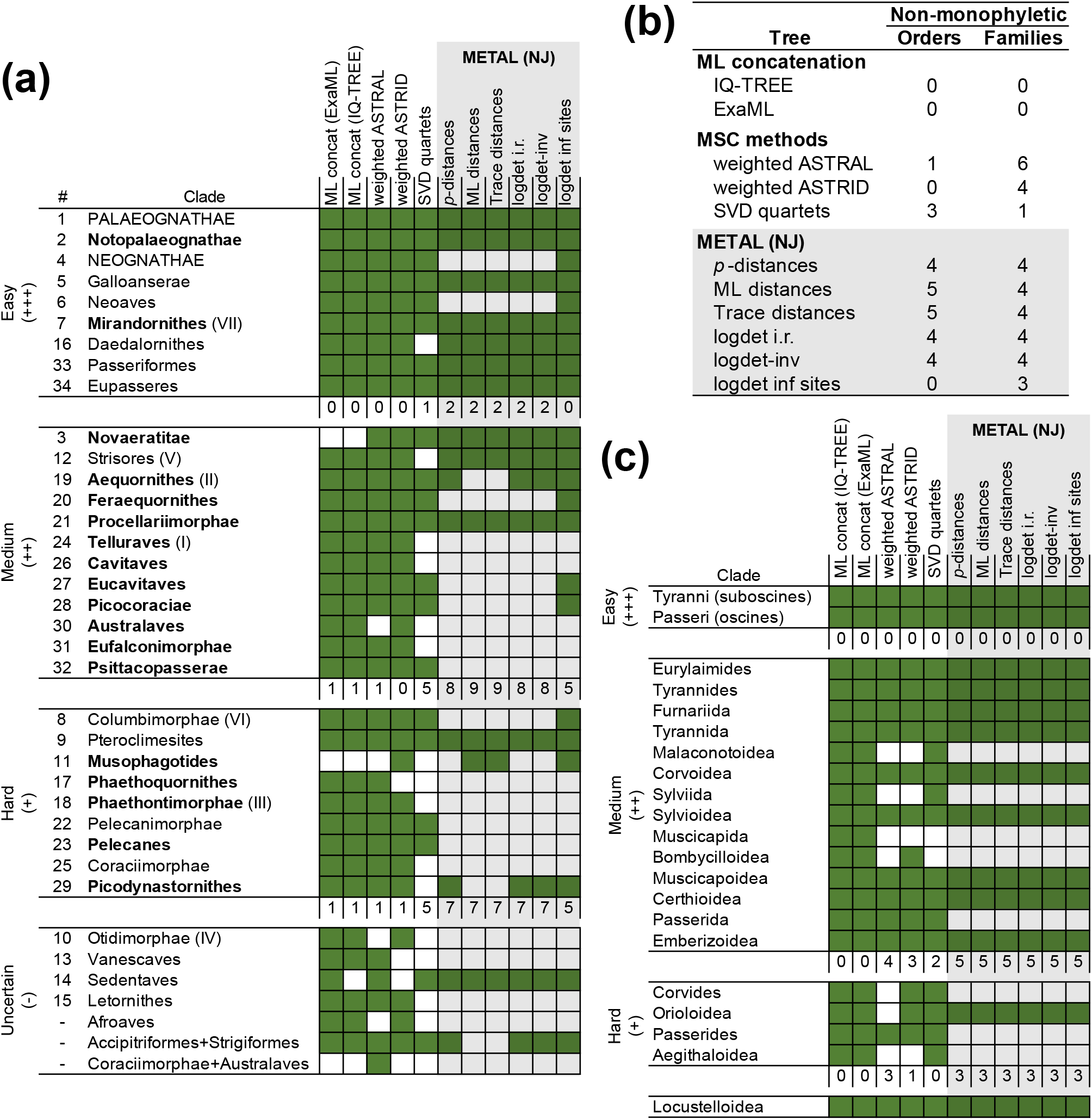
Recovery of clades in analyses of the complete allfam dataset. Grids show the presence (green) or absence (white) of clades in trees generated by each method. Reliable clades were divided into three groups (easy, medium, and hard); the fourth group is clades that were not viewed as reliable. Numbers of reliable clades that were not recovered as monophyletic groups are listed below each set of clades. (**a**) Recovery of deep clades (primarily supraordinal, but also including the taxon-rich clades Passeriformes and Eupasseres). (**b**) Numbers of non-monophyletic orders and families. For this dataset we were able to assess the monophyly of 39 orders and 93 families. We did not include the IOC family Tityridae (which is not reliable) in the count presented. (**c**) Recovery of suprafamilial passerine clades. The model for ML and trace distances was GTR+I+Γ. METAL analyses are shaded gray. All trees are available in the File S1.

The performance of the MSC methods based on the “reliable clade recovery” criterion was not as good as ML concatenation (Fig. 3). The best MSC method was wASTRID, followed closely by wASTRAL. Both gene tree summary methods recovered many deep branches correctly (Fig. 3a), although the weighted ASTRID tree had limited support (see File S1). Most problems for wASTRID and wASTRAL were closer to the tips of the tree (Fig. 3b,c).

SVDquartets performed worse than ML concatenation or the gene tree summary methods, though the problems with SVDquartets were different. Deep in the tree, SVDquartets failed to recover one easy supraordinal clade, five medium supraordinal clades, and five hard supraordinal clades. It also failed to recover three orders (Charadriiformes, Apodiformes, and Gruiformes). None of those orders were hard clades; they were present in 66.7%, 74.1%, and 83.3% of the individual gene trees from (Reddy et al., 2017), respectively (see Table S4). SVDquartets performed better than wASTRAL and wASTRID closer to the tips of the tree, recovering more superfamilies and families than either gene tree summary method (Fig. 3b,c), but it never performed as well as ML concatenation.

METAL had the worst performance of all methods tested, with the NJ trees for all distance estimators failing to recover large numbers of reliable clades at all depths in the tree (Fig. 3). The best performing METAL analyses was NJ of logdet inf sites distances; it recovered more reliable supraordinal clades than any other METAL analysis (Fig. 3a) and it was the only METAL analysis that supported monophyly of all orders (Fig. 3b). Many rearrangements relative to expectation had high support in the METAL trees (in some cases unexpected rearrangements had 100% support; see File S1).

### Missing data affects branch length estimates

In previous large-scale studies of avian phylogeny (Hackett et al., 2008; Berv and Field, 2018; Braun et al., 2019; Braun and Kimball, 2021; Ji et al., 2022) certain taxa routinely display long terminal branches in analyses using a variety of data types (we have highlighted several examples of these lineages with with bird silhouettes and black labels in Fig. 4). However, several unexpected taxa with exceptionally long branches were also evident in the ML concatenation trees generated using the complete allfam dataset. All of these unexpected long branch taxa also had poor sequence recovery (red labels in Fig. 4). Despite the long branches, none of the taxa with poor sequence recovery appeared to be misplaced in the ML concatenation tree based upon expectations from prior studies. For example, *Moho nobilis* (the extinct Hawaii Oo; Passeriformes: Mohoidae) and *Hypocolius ampelinus* (Grey Hypocolius; Passeriformes: Hypocoliidae) were placed within Bombycilloidea, as expected based on analyses of individual gene regions (Fleischer et al., 2008; Spellman et al., 2008). The taxon with the worst sequence recovery, *Strigops habroptila* (Kakapo; Psittaciformes: Nestoridae) provided an even more striking example. Although *Strigops* had a long terminal branch it was placed sister to the confamilial, *Nestor notabilis* (Kea); both *Strigops* and *Nestor* are parrots endemic to New Zealand.

**Figure 4.**
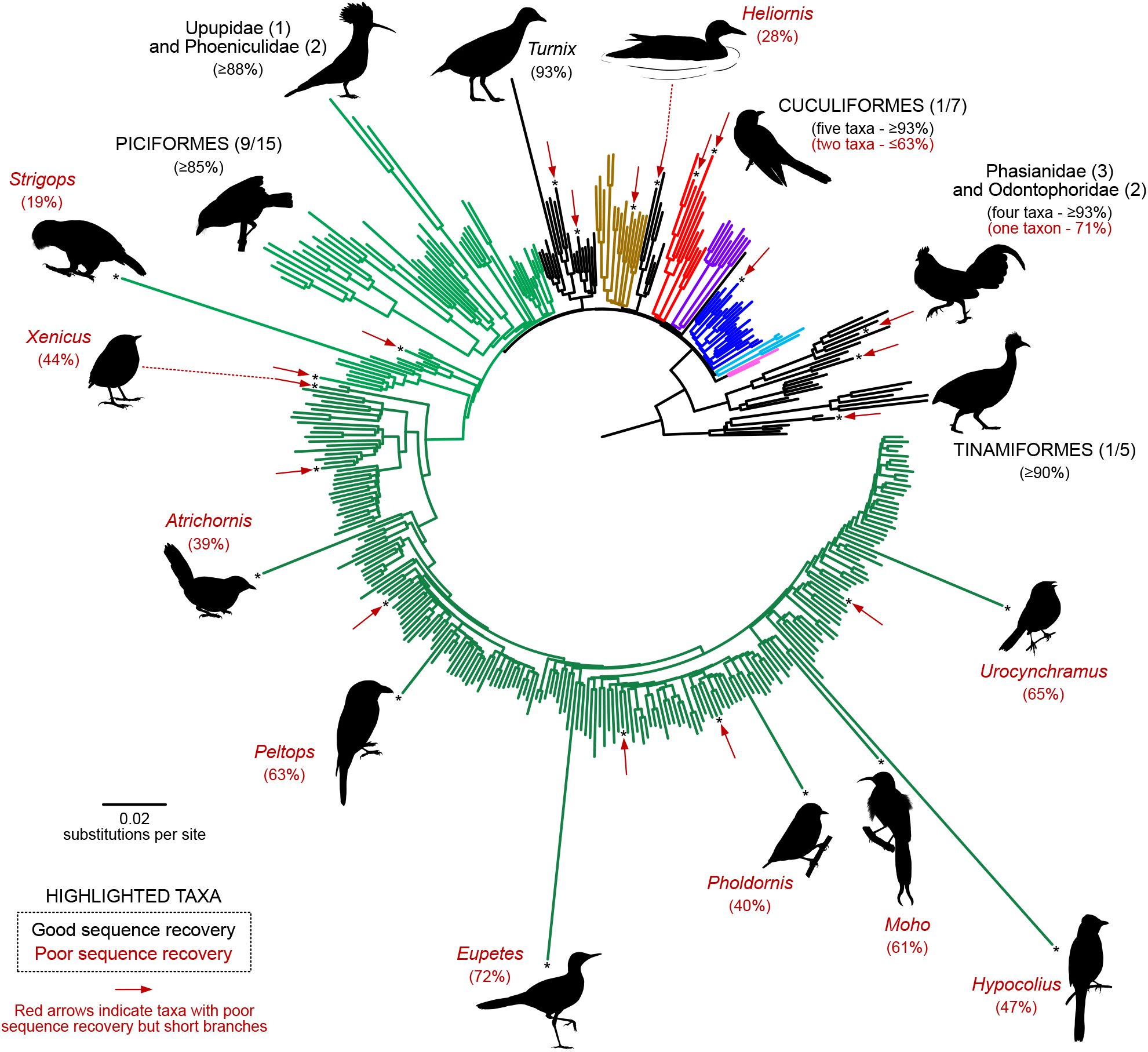
Branch length heterogeneity in the ML concatenation tree reflects a combination of biological signal and methodological artifacts. The IQ-TREE topology for the complete allfam dataset presented as a circular phylogram. With the exception of Passeriformes, the colors match those in the other figures; Passeriformes are presented in a darker shade of green than the other Telluraves to emphasize the extensive sampling within that order. Taxa with long branches are highlighted with silhouettes. Most taxa with long branches are in clades that appear to have accelerated sequence evolution (emphasized with silhouettes and black labels). The number of species sampled in each of those clades is indicated in parentheses, presented either as the number of sampled families and species (for orders) or as the number of species (for families). A small number of taxa with isolated long branches had poor data recovery (emphasized with silhouettes and red labels); these long branches are likely to be artifactual. Data recovery is indicated for all taxa with silhouettes; recovery is measured as the percentage of resolved bases in the final alignment relative to the taxon with the maximum number of resolved bases (2,012,920 bp). All taxa with *<*75% resolved bases are indicated with asterisks at the tips of branches and, when the branch lengths are not anomalous, they are indicated with short brick red arrows. Bird silhouettes were drawn by Min Zhao.

The majority of taxa with poor sequence recovery did not have especially long branches (only nine of the 27 taxa with poor sequence recovery had noticeably long branches). We have highlighted two poorly-sampled taxa with relatively short branches in Fig. 4, the passerine *Xenicus gilviventris* (New Zealand Rockwren) and the gruiform *Heliornis fulica* (Sungrebe), to emphasize that these poorly-sampled taxa with relatively short branches may have branch lengths that are either slightly longer than their sister or even slightly shorter. Some of these poor sequence recovery taxa (all of which are indicated with short red arrows and asterisks at the ends of branches in Fig. 4) are in lineages that have long branches in other studies.

For example, most Cuculiformes that we sampled had good sequence recovery and they have long branches, as we expected based on other studies. However, two Cuculiformes (*Centropus viridis* [Philippine Coucal; Cuculiformes: Cuculidae] and *Phaenicophaeus curvirostris* [Chestnut-breasted Malkoha; Cuculiformes: Cuculidae]) had poor sequence recovery. Although those two poorly-sampled Cuculiformes had long branches, they did not appear anomalous relative to the well-sampled Cuculiformes. None of the taxa with poor sequence recovery but short branches appeared to be misplaced in the ML concatenation tree.

Taxa with poor sequence recovery appeared to be problematic in many MSC analyses. This was especially true for METAL; NJ of most distance metrics placed *Strigops* sister to all other birds rather than with other parrots (File S1). SVDquartets was the exception to this pattern; the cases of ordinal non-monophyly that we observed in the SVDquartets tree based on the complete dataset did not have an obvious association with missing data (File S1). In fact, SVDquartets recovered a *Strigops*+*Nestor* clade in analyses of the complete allfam dataset, suggesting that it is relatively robust to missing data, at least for relationships closer to the tips of the tree. Nevertheless, the poor performance of the gene tree summary methods and METAL when used to analyze the complete allfam dataset indicate that the impact of missing data make it difficult to assess overall method performance.

### Reducing missing data improves the performance of some MSC analyses

We excluded the 27 taxa with the worst sequence recovery (*<*75% resolved bases in the final alignment) to examine method performance using a dataset with less missing data (the excluded taxa are listed in Table S3). Given the high degree of similarity between the ML concatenation trees generated using IQ-TREE and ExaML (Fig. 2 and supplementary), we limited our ML concatenation analysis of this “well-sampled” allfam dataset to IQ-TREE, which was faster. This resulted in a tree that was very similar to the ML concatenation trees for the complete dataset (compare ML concatenation trees in File S1 to those in File S2). The impact of taxon exclusion on MSC approaches was more variable (Fig. 5 and File S2). Recovery of supraordinal clades was slightly worse for the gene tree summary methods when we used the well-sampled allfam dataset; relative to the analyses using the complete dataset, wASTRAL failed to recover three additional reliable clades and wASTRID failed to recover one additional reliable clade. Unsurprisingly, the additional clades that those methods failed to recover were in our hard category (e.g., compare Figs. 3a and 5a). In contrast, the gene tree summary methods generally had better recovery of more derived clades, like orders, superfamilies, and families (Fig. 5b,c). Deleting taxa with poor sequence recovery had a more universally positive impact on METAL analyses (compare Figs. 3 and 5), although even the best METAL analysis (NJ with logdet inf sites distances) still recovered fewer reliable clades than ML concatenation.

**Figure 5.**
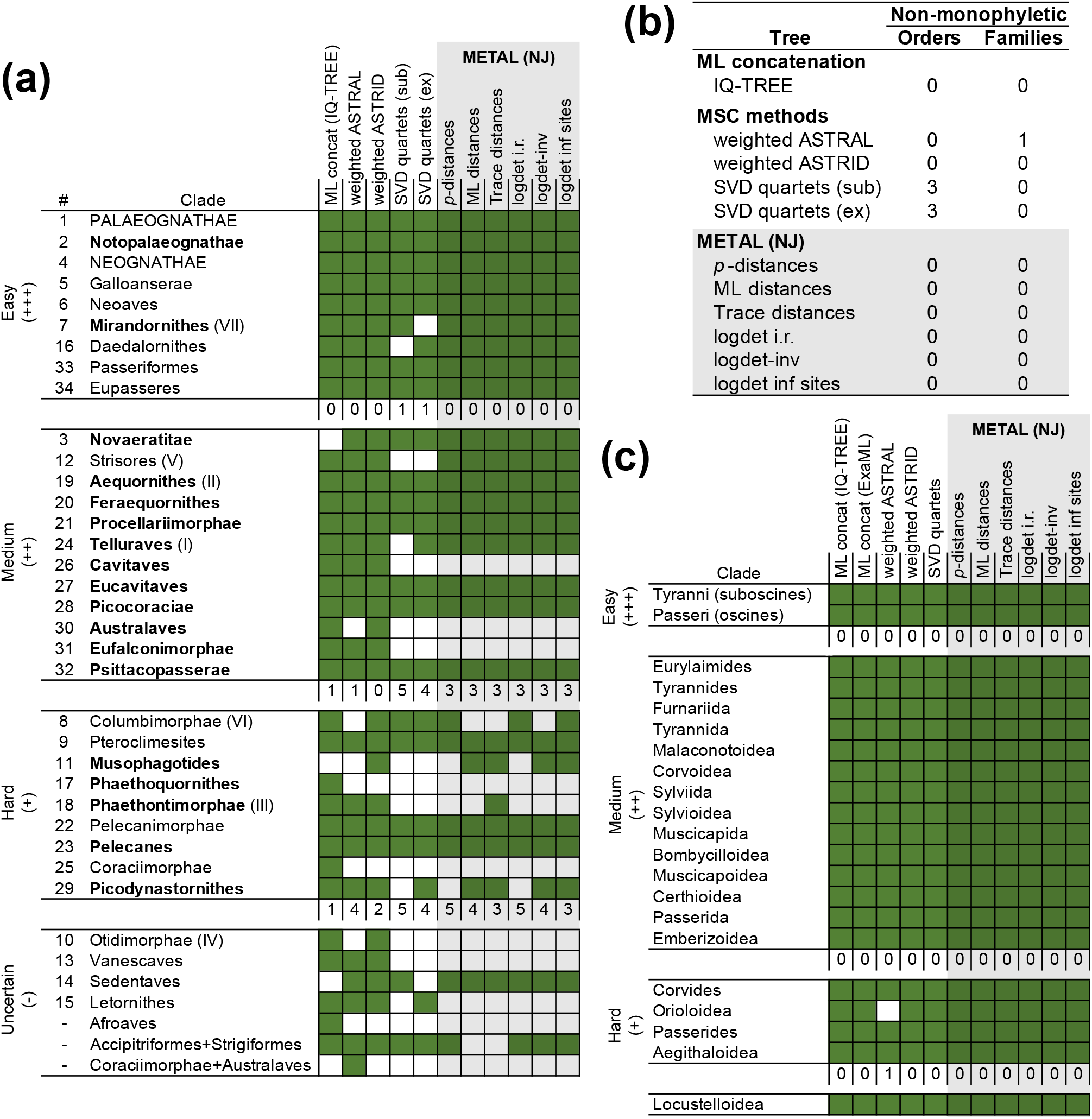
Recovery of clades in analyses of the well-sampled allfam dataset. Grids show the presence (green) or absence (white) of clades in trees generated by each method; clades were divided into groups in a manner identical to Fig. 3. Two SVDquartets analyses are included, one with subsampling (sub) and one with exhaustive sampling within subtrees (ex) (**a**) Recovery of supraordinal clades (along with Passeriformes and Eupasseres). (**b**) Numbers of non-monophyletic orders and families. For this dataset we were able to assess the monophyly of 39 orders and 83 families. (c) Recovery of suprafamilial passerine clades. The model for ML and trace distances was GTR+I+Γ. METAL analyses are shaded gray. All trees are available in File S2.

The performance of SVDquartets was still poor based on the reliable clade criterion, but its behavior was different from the other methods. SVDquartets did not appear especially sensitive to missing data. However, it failed to recover some easy ordinal and supraordinal clades (e.g., Charadriiformes, Apodiformes, and Gruiformes; see above). One limitation of SVDquartets is the need to evaluate a large number of quartets when the taxon sample is large, which can be computationally demanding. We initially addressed this limitation by sampling 10^7^ quartets (*∼*1% of the total distinct quartets). Obviously, SVDquartets might exhibit better behavior if we sampled quartets exhaustively. To accomplish this and give SVDquartets the best opportunity to compete with other methods we divided the well-sampled allfam taxon set into two uncontroversial subsets (chiefly passerines and non-passerines, with the former subset including six non-passerine outgroups and the latter including six passerines to place the order) and generated trees by sampling quartets exhaustively within each subset. However, that strategy did not result in a substantive improvement in the behavior of SVDquartets. The SVDquartets exhaustive sampling tree did address the issues of Charadriiformes and Apodiformes monophyly, but Gruiformes was still nonmonophyletic (it was split into three distinct groups) and one additional order became non-monophyletic (Musophagiformes, an order that was supported in all of the individual gene trees from Reddy et al., 2017). The observation that the recovery of reliable clades varies between the random subsampling and exhaustive analyses for subsets suggests that different random subsamples might exhibit different behaviors; this variability might also emerge if the taxa are subdivided for exhaustive sampling in different ways. Regardless of the details, this behavior is undesirable. SVDquartets was the only method that failed to recover two of the easy supraordinal clades (Mirandornithes with exhaustive sampling of non-passerines and Daedalornithes with random subsampling; see Fig. 5a).

All METAL trees shared a number of features that suggest their poor performance relative to other methods might reflect long-branch attraction. Telluraves provides an excellent illustration of the evidence for long-branch attraction (Fig. 6). The METAL trees split Telluraves into two clusters (Fig. 6a), one characterized by taxa with long-branches and a second by short-branch taxa (the short-branch lineages comprise the four orders of raptorial birds and the monotypic order Leptosomiformes, which we call “raptors+”). In sharp contrast, the ML concatenation tree shows three separate clusters of long-branch (emphasized with light purple boxes in Fig. 6b) separated by the five raptor+ orders, which all have relatively short branches. This interspersion of longand short-branch taxa in the ML concatenation topology is consistent with three strongly corroborated clades (black dots in Fig. 6b) that also have support in analyses of TE insertions and indels (Suh et al., 2015; Houde et al., 2019; Gatesy and Springer, 2022). Moreover, the wASTRAL and wASTRID trees also separate the long branch lineages into three clusters (File S2), so intermixing of long- and short-branch taxa is not a feature exclusive to ML concatenation.

**Figure 6.**
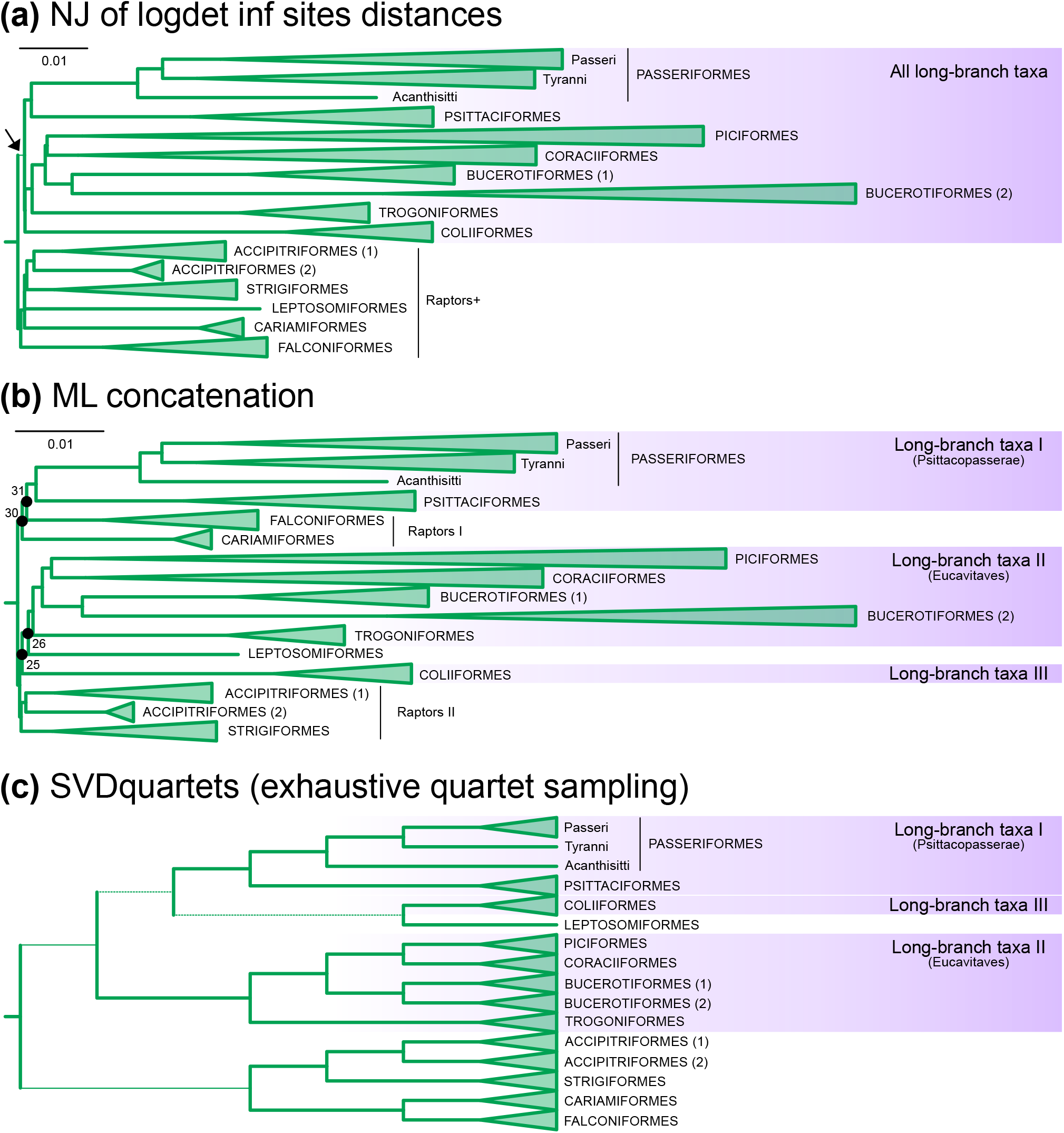
Clustering of long-branch taxa in METAL and SVDquartets trees. This figure is limited to Telluraves (clade 24), where the clustering of long branch taxa was most obvious (note the color). (**a**) METAL tree (NJ of logdet inf sites distances). The branch uniting the long branch taxa had low (54%) bootstrap support (indicated with a thin branch and arrow); the remaining branches all had *>*90% bootstrap support. (**b**) ML concatenation. Numbers adjacent to black dots on the ML concatenation tree are clade identifiers (see Fig. 1). All branches had *>*90% bootstrap support. (**c**) SVDquartets tree (exhaustive quartet sampling of the non-passerine taxon sample). This tree is presented as a cladogram because SVDquartets does not provide branch length estimates. Thin branches have *<*90% bootstrap support and dashed branches have *<*50% support. Most taxa in these trees are grouped into orders; the length of each clade triangle corresponds to the length of the longest branch in that order. Three orders were subdivided to better present branch length heterogeneity within those orders. The three major passerine subclades are defined in Fig. 1b. Bucerotiformes (1) comprises Bucerotidae and Bucorvidae (hornbills) and Bucerotiformes (2) comprises Upupidae and Phoeniculidae (hoopoes, woodhoopoes, and scimitarbills). Accipitriformes (1) comprises Accipitridae, Pandionidae, and Sagittariidae (hawks, eagles, Old World vultures, Osprey, and Secretarybird) and Accipitriformes (2) comprises Cathartidae (New World vultures). Trees with support values are available in File S2.

There were several additional differences between the METAL trees and trees based on ML concatenation and wASTRAL/wASTRID that are consistent with the hypothesis that METAL is highly susceptible to long-branch attraction. The SVDquartets Telluraves topology resembles the METAL topology (Fig. 6c), suggesting that the method may be at least somewhat sensitive to long-branch attraction. However, the support was lower overall (note the thin branches in Fig. 6c) and Leptosomiformes was placed inside the long-branch cluster rather than within a raptors+ cluster.

We focused on the presence or absence of clades in the optimal trees, but some conflicts are poorly supported. For example, collapsing branches with low support in the SVDquartets tree would yield an unresolved topology for Telluraves (Fig. 6c), eliminating the conflicts with reliable clades. To examine this issue in more detail, we generated a set of partially resolved trees where branches with *<*95% support were collapsed. All collapsed trees were relatively well-resolved (Table 4), with the ML concatenation tree being the best-resolved. The collapsed wASTRID and SVDquartets (exhaustive quartet sampling) trees had fewest conflicts with the ML concatenation tree. However, collapsing branches with *<*95% support did not eliminate the raptors+ clade in the logdet inf sites METAL tree (that unexpected clade had 100% support).

**Table 4.** Nodes with *≥*95% support in analyses of the well-sampled allfam dataset (369 taxa total)

| Tree | Resolved Branches | Conflicting Branches<sup>1</sup> |
| --- | --- | --- |
| ML concatenation | 347 (94.8%) | — |
| Standard MSC methods |  |  |
| wASTRAL | 317 (86.6%) | 17 (16) |
| wASTRID | 292 (79.8%) | 9 (7) |
| SVDquartets | 304 (83.1%) | 6 (4) |
| METAL trees |  |  |
| *p*-distances | 315 (86.1%) | 30 (28) |
| Trace distances | 322 (88.0%) | 27 (24) |
| ML distances | 323 (88.3%) | 26 (23) |
| logdet i.r. distances | 317 (86.6%) | 31 (29) |
| logdet-inv distances | 315 (86.1%) | 28 (25) |
| logdet inf sites distances | 323 (88.3%) | 18 (16) |

<sup>1</sup> Number of branches in each tree that conflict with the ML concatenation tree. The parenthetical is the number of branches that conflict with branches in the ML concatenation tree that have $\geq 95\%$ support.

### Does more data improve the performance of METAL (or other phylogenomic methods)?

Judged by the reliable clade recovery criterion, all MSC methods failed to perform as well as ML concatenation on the allfam datasets. This was especially true for METAL, which exhibited the worst performance of all methods tested. This apparent poor behavior of METAL could reflect bias (e.g., greater sensitivity to long-branch attraction) or it could reflect the use of insufficient data for METAL to perform well. Analyzing a larger dataset should yield trees similar to those we obtained using the allfam dataset, likely with higher support, if the problem is bias. On the other hand, if METAL requires more data, then a larger dataset should improve its performance.

The Jarvis UCE and intron datasets, had more sites than the allfam datasets (4.5x and 9.4x, respectively; Table 2), but only 48 taxa. This reduced the number of reliable clades that could be assessed to 26 (Table 3). The additional sites in the Jarvis UCE data appears to have improved the recovery of reliable clades for wASTRAL and wASTRID (Fig. 7). Both gene tree summary methods recovered all reliable clades that could be assessed using this taxon sample; in contrast, similar analyses using the well-sampled allfam dataset failed to recover five (wASTRAL) or two (wASTRID) of the reliable clades that could be assessed using the Jarvis taxon sample. METAL may have exhibited a modest improvement, at least when logdet inf sites distances and (to a lesser degree) when ML and trace distances were used (Fig. 7). NJ of logdet inf sites recovered three of the four supraordinal clades in Telluraves that group long- and short-branch taxa (purple shading in Fig. 7) and the fourth was not placed in a distinct position with high support (File S3). The use of other distance estimators recovered fewer of those putative long-branch attraction “indicator clades” and some of the relevant taxa had high (*≥*95%) support for their unexpected placements (File S3). The SVDquartets analysis did not exhibit a substantive improvement (we do not view the shift from five to four non-monophyletic hard clades as strong evidence of an improvement). The poor behavior of SVDquartets did not reflect incomplete quartet sampling because the limited number of taxa in the Jarvis datasets allowed us to use exhaustive quartet sampling.

**Figure 7.**
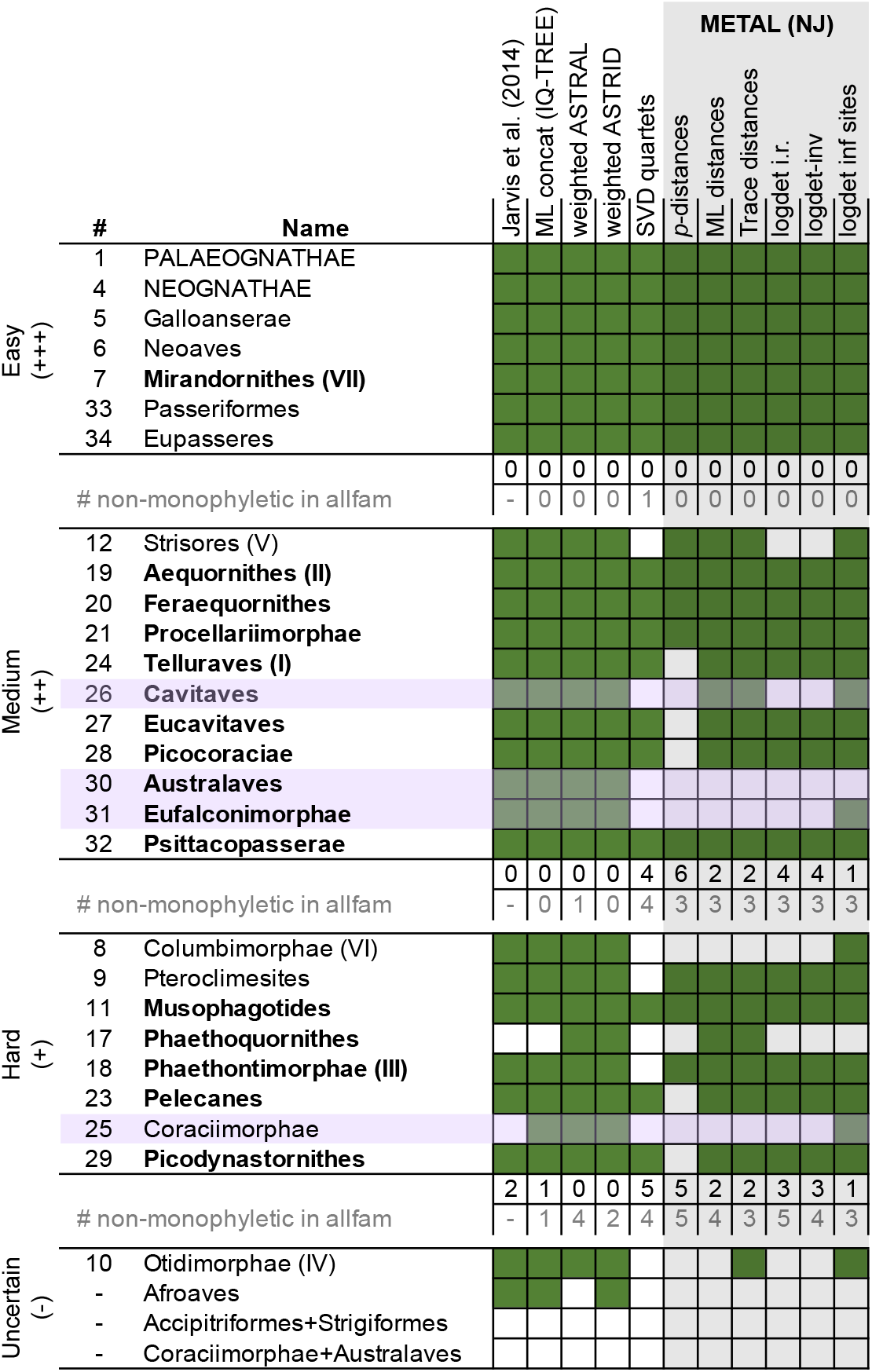
Recovery of clades in analyses of the Jarvis UCE dataset. Grids show the presence (green) or absence (white) of clades. Numbers of reliable clades that were not recovered are listed immediately below each set of clades in black text and the numbers of reliable clades that were not recovered as monophyletic groups in comparable analyses of the well-sampled allfam dataset are listed below those values in gray text (these numbers differ from Fig. 5 because they are limited to clades that can be assessed using the Jarvis taxon sample). METAL analyses are shaded gray; clades in Telluraves that comprise both longand short-branch taxa are emphasized (also see Fig. 6) in purple. The model for ML and trace distances was GTR+I+Γ. We have included the UCE tree from Jarvis et al. (2014), which was estimated using RAxML v. 7.3.5 (Stamatakis, 2006) without TAPER masking; that tree had a small topological difference from our ML concatenation tree. All of the Jarvis UCE trees estimated for this study are available in File S3.

The Jarvis intron data has almost threefold more informative sites than the Jarvis UCE dataset (Table 2), so if more data are critical to the performance of METAL, analyses of the intron data should provide the best results. Unfortunately, the number of reliable clades that remained problematic in analyses of Jarvis UCE data is limited. Both gene tree summary methods exhibited good behavior and METAL with the best distance metrics only had problems with Telluraves clades that include long- and short-branch taxa (Fig. 7). Thus, we expect any improvement in the recovery of reliable clades to be limited to those clades (it is also possible that some idiosyncratic aspect of intron evolution will cause other nodes to become problematic in specific analyses, but that seems unlikely).

ML concatenation and the gene tree summary methods had equally good behavior when used to analyze the Jarvis intron data, with each method only failing to recover a single hard clade (Fig. 8a). SVDquartets performed poorly, just as it did with the Jarvis UCE data. METAL analyses of the intron data using ML and trace distances with the GTR+Γ model recovered all of the reliable clades in Telluraves that include both short-branch and long-branches (Fig. 8a). The best distance estimator in analyses of the Jarvis UCE data and the allfam UCE data was always logdet inf sites; analyses using that distance estimator was only slightly worse; the only indicator clade that NJ of logdet inf sites distances failed to recover was Australaves (clade 30). These results suggest that the Jarvis intron dataset provides sufficient information for relatively good performance of METAL (at least with some distance estimators).

**Figure 8.**
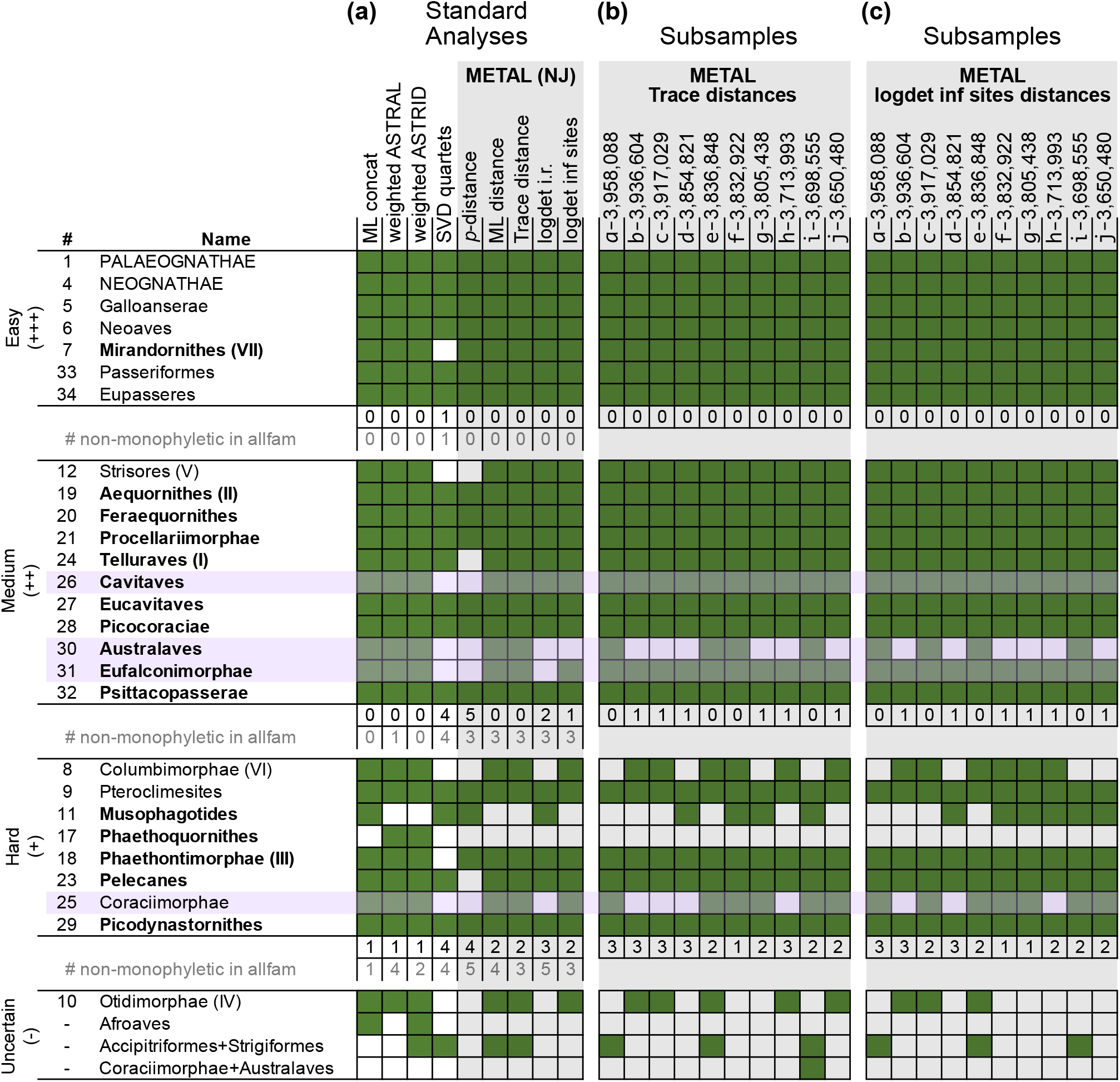
Recovery of clades in analyses of the Jarvis intron dataset. Grids show the presence (green) or absence (white) of clades. (**a**) Results of the set of “standard analyses” that we used with all datasets. The Ml concatenation tree had a topology identical to the intron ML tree in Jarvis et al. (2014). The model for ML and trace distances was GTR+Γ. We have omitted NJ of logdet-inv distances because the best-fitting model for the Jarvis intron data did not include invariant sites. (**b**) METAL analyses using NJ of trace distances (GTR+Γ with *α*=2) to analyze subsets of the Jarvis intron data. Each subset comprises 900 randomly selected locus alignments. Numbers of informative sites are listed after each subset identifier (e.g., subset ‘a’ has 3,958,088 informative sites). The number of informative sites in the Jarvis UCE dataset falls between subsets ‘d’ and ‘e.’ (**c**) METAL analyses using NJ of logdet inf sites distances to analyze subsets of the Jarvis intron data. METAL analyses are shaded gray in all panels and the long-branch attraction indicator clades are highlighted with light purple. All trees are available in File S3.

METAL with trace and ML distances failed to recover two clades (Fig. 8a), both of which warrant mention. First, neither ML concatenation nor METAL recovered Phaethoquornithes, although the conflicts with that clade differ between the methods. METAL trees had a Phaethontimorphae+Opisthocomiformes clade (node *α* in Fig. 9a) regardless of the distance estimator used. The unexpected Phaethontimorphae+Opisthocomiformes was sister to Aequornithes, so the rearrangement relative to Phaethoquornithes is actually the movement of Opisthocomiformes, a monotypic order that moves to a position sister to Phaethontimorphae. This unexpected Phaethontimorphae+Opisthocomiformes is strongly supported in all METAL analyses. ML concatenation failed to recover Phaethoquornithes because it placed Phaethontimorphae sister to Telluraves (node *α* in Fig. 9b). We expected the ML concatenation tree for the Jarvis intron to have the Phaethontimorphae+Telluraves clade because the ML concatenation trees for the intronic data in Jarvis et al. (2014) had the same clade. Indeed, Reddy et al. (2017) highlighted the position of Phaethontimorphae as a potential data type effect (where the Phaethontimorphae+Telluraves clade was supported non-coding data). This study emphasizes that analyses of noncoding data do not consistently support Phaethontimorphae+Telluraves. Both ML concatenation and gene tree summary methods support Phaethoquornithes in analyses of the allfam datasets (Figs. 3 and 5). Gene tree summary analyses using both Jarvis datasets also supported Phaethoquornithes (Figs. 7 and 8a). When those results are considered in light of the support for Phaethoquornithes in analyses of TE and indel data (Suh et al., 2015; Houde et al., 2019) they corroborate the classification of Phaethoquornithes as a hard but reliable clade.

**Figure 9.**
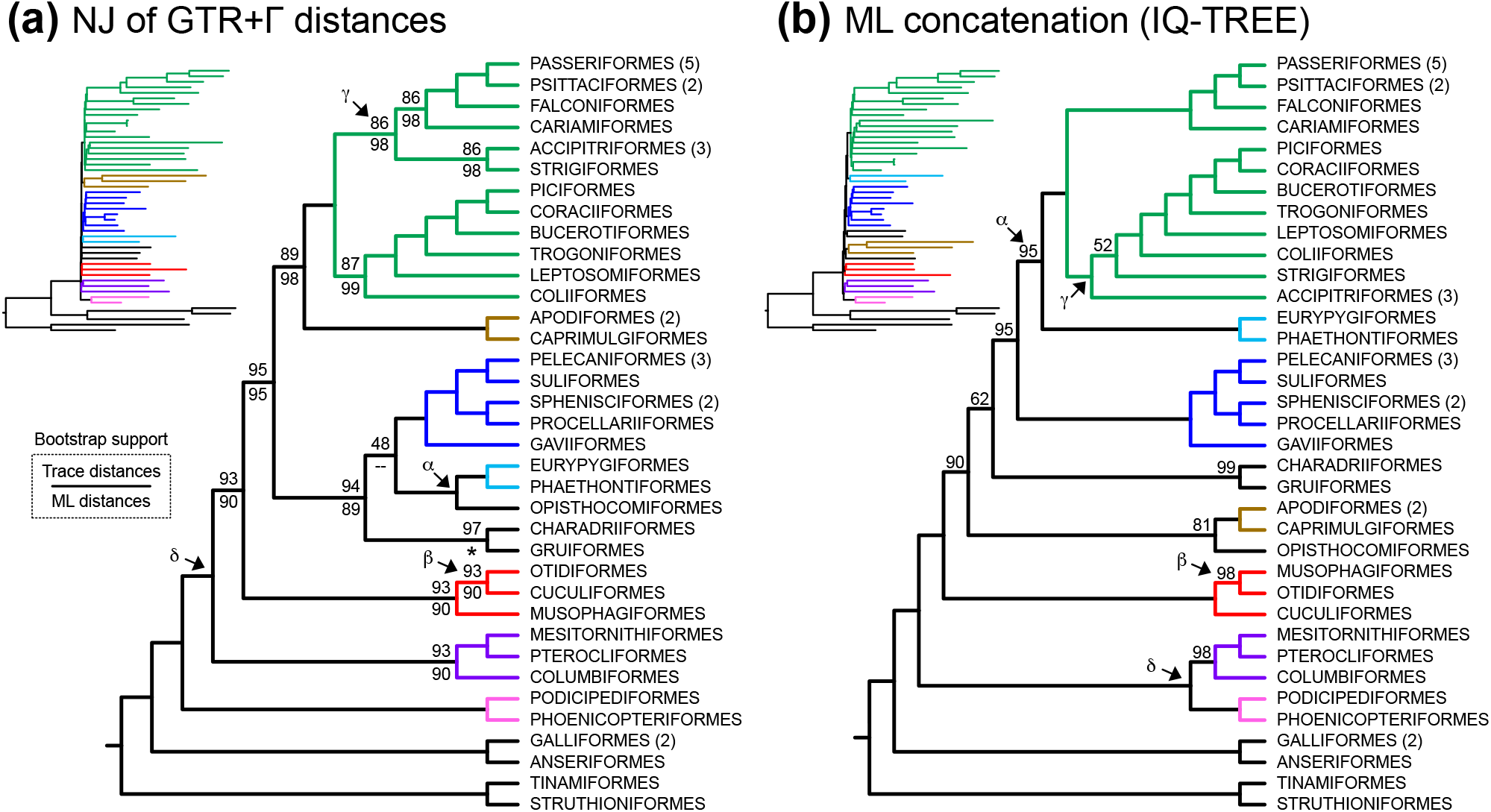
Estimates of avian ordinal relationships based on the Jarvis intron dataset. Cladograms showing relationships among orders. The number of sampled taxa from each order is listed parentheses; absence of a parenthetical number indicates that a single taxon was sampled. Phylograms are presented as insets to illustrate branch lengths. (**a**) METAL tree for the Jarvis intron data. The topology reflects trace distances using the GTR+Γ model. Bootstrap support values above branches are based on trace distance and those below branches are based on ML distances. Branches with complete (100%) support are unlabeled. There is one topological difference between the two versions of the GTR+Γ distances (note the ‘–’ for ML distance support) and one node with 97% support based on trace distances but complete (100%) support based on ML distances (note the asterisk). (**b**) ML concatenation tree for the Jarvis intron data. Nodes indicated with Greek letters are described in the text. Trees with all taxa are available in File S3.

The second clade of interest is Musophagotides, which we also classified as a hard but reliable clade. However, METAL analyses using the best distance estimators failed to recover Musophagotides, instead placing Cuculiformes sister to Otidiformes (node *β* in Fig. 9a), a topology that matches the ML concatenation tree for the allfam UCE dataset (Fig. 2). This alternative topology is a local rearrangement (note the positions of node *β* in Fig. 9a vs Fig. 9b) and the gene tree summary methods both recover the same topology as METAL with the best distance estimators (Supplementary Fig. S2). Although that result might suggest that the conflict is between ML concatenation and MSC methods conflict, we note that all analyses of the Jarvis UCE data recover Musophagotides; when that result is considered in light of the observation that analyses of TE and indel data also support Musophagotides (Suh et al., 2015; Houde et al., 2019) it seems reasonable to retain the Musophagotides as a hard but reliable clade. Cuculiformes is a long branch clade (see Fig. 4); METAL might be problematic due to shifts in evolutionary rates in this case.

The role of data type (intron versus UCE) in these analyses is unclear, although there are hints that it might play a role (e.g., based on the conflict regarding Musophagotides). Indeed, our assertion that the recovery of all long-branch attraction indicator clades (light purple highlights in Fig. 8) provides evidence that METAL exhibits improved behavior when used with the Jarvis intron data might reflect either dataset size or data type. If the issue is primarily dataset size, then randomly selected subsets of the Jarvis intron data that are similar in size to the Jarvis UCE data (where size is measured in numbers of parsimony informative sites) would be expected to exhibit behavior similar to analyses of the UCE data. This appeared to be the case; we observed long-branch attraction indicator clades Australaves and Coraciimorphae were only recovered in METAL analyses of some random subsets (Fig. 8b). This indicates that the failure to recover one of those clades in the METAL analysis of the Jarvis UCE data can be explained by dataset size, although it does not exclude the possibility of a data type effect combined with a dataset size effect. In fact, the behavior of METAL analyses of the Jarvis intron and UCE datasets suggests an impact of data type (compare recovery of the indicator clades in Fig. 7 and Fig. 8a); specifically, NJ of ML and trace distances appear to exhibit better behavior than logdet inf sites distances with the intron data. On the other hand, the number of reliable clades with variable recovery is limited, so it is not absolutely clear that NJ of logdet inf sites distances exhibits worse behavior. Indeed, all indicator clades were recovered in analyses using four out of ten subsamples when logdet inf sites distances were used (Fig. 8c), a result identical to NJ with trace distances. Thus, the hypothesis that there is a data type effect remains plausible, but it is subtle if it exists and it is not corroborated at this point.

## Discussion

The goal of this study was to compare the behaviors of various methods to estimate species trees using a novel avian dataset that included representatives of almost all avian families. We included METAL—defined broadly as the use of distance analyses with phylogenomic data for species tree estimation—as one of the MSC methods. Using a large number of reliable clades permitted meaningful evaluation of different methods. This method to evaluate trees revealed that ML concatenation exhibited good performance, yielding trees that included most reliable clades regardless of whether or not taxa with poor data recovery were included. Other methods generally performed worse than ML concatenation, although gene tree summary methods generally exhibited better performance than MSC methods that use concatenated data (SVDquartets and METAL). To determine whether the relatively poor performance of MSC methods reflected the number of sites or type of data in the allfam datasets, we used the same methods to reanalyze two published datasets with more sites but fewer taxa. Analyses of the larger datasets provided evidence that almost all of the methods we examined can exhibit good performance when sufficient data are available.

### The impact of missing data on phylogenomic analyses is complex

Although ML concatenation appeared to yield accurate estimates of tree topology, it produced poor estimates of branch lengths for taxa with large amounts of missing data. Large amounts of missing data are known to create problems for branch length estimation (Darriba et al., 2016) and it has been noted in other UCE studies where missing data are a problem (Kimball et al., 2021; Nash et al., 2024). However, the number of taxa with problematic branch lengths was limited and it did not have an obvious impact on topological accuracy. In contrast, missing data appeared to have a greater negative impact on topological estimation with the other methods we examined, although there was some complexity in those results. First, removing poorly-sampled taxa and using gene tree summary methods (wASTRAL and wASTRID) appeared to result in slightly worse behavior deep in the tree but better behavior closer to the tips of the tree. SVDquartets exhibited different behavior; it recovered more families and superfamiles than either of the gene tree summary methods in (Figs. 3b,c and 5b,c) but it was more problematic at the level of orders and supraordinal clades regardless of the inclusion or exclusion of taxa with poor sequence recovery. METAL was most sensitive to missing data: it behaved fairly poorly overall when it was used with the complete allfam UCE dataset (Fig. 3). However, METAL recovered all reliable clades below the supraordinal level once poorly sampled taxa were removed, regardless of which distance estimator was used (Fig. 5b,c).

All methods exhibited fairly good behavior with the larger Jarvis datasets. This may seem surprising since those datasets have a larger proportion of gaps and missing data than the allfam datasets (Table 2). However, the complete allfam dataset has two issues with the potential to exacerbate the negative impact of missing data. First, the variance among taxa in the amount of missing data is extreme, ranging from less than 2% of sites in some taxa to more than 80% in the taxon with the worst sequence recovery. Second, the distribution of missing data in the complete allfam dataset is likely to be problematic because poor sequence recovery disproportionately affects sites that are further from the conserved core of the UCE locus that hybridizes with the probes (Hosner et al., 2015). Thus, the missing data for taxa with poor sequence recovery primarily lack the more variable flanking sequences. Those flanking sequences are the source of most phylogenetic signal in UCE datasets. Both Jarvis datasets were extracted from whole genomes, so the missing data are more evenly distributed throughout the sequences.

Non-independence of the underlying models for nucleotide and indel evolution can lead to inconsistency for distance methods when gaps are treated as missing data. McTavish et al. (2015) presented a proof showing that “…if the indel process and substitution process are independent, treating gaps as missing data will not cause statistical inconsistency of distance-based tree inference.” However, the conditions assumed in that proof are violated if a subset of sites are both free of substitutions and indels. The technical issues introduced by sequence capture mean that the distribution of missing data in our alignments are not independent of the substitution model because the ultraconserved regions that hybridize with the sequence capture probes will be captured even when the informative flanking sequences are not captured. This means that missing data will disproportionately be associated with more rapidly evolving sites, a situation analogous to the one McTavish et al. (2015) found to be potentially problematic. In contrast, the distribution of gaps and missing data in the Jarvis datasets are more independent of the patterns of sequence evolution, and therefore less problematic for METAL.

### Differential performance of ML concatenation and MSC methods

MSC methods were developed to address the bias in ML concatenation (Edwards, 2009; Mirarab et al., 2021); clades that are present in trees generated using most (or all) MSC methods but absent in the ML concatenation tree would provide empirical evidence for that bias. Novaeratitae (clade 3) was the only example of such a clade that we found; Apterygiformes is a deep branch within Palaeognathae in ML concatenation analyses (Files S1 and S2) but sister to Casuariiformes (i.e., Novaeratitae is present) in all MSC trees (Figs. 3a and 5a, also see Files S1 and S2). Consistent with our results, Apterygiformes and Casuariiformes are separated in a number of published ML concatenation trees for phylogenomic datasets (Prum et al., 2015; Braun and Kimball, 2021; Kuhl et al., 2021) and an MSC analyses using those the Prum et al. (2015) data included Novaeratitae (compare the concatenated analyses and the ASTRAL analysis of RY-coded data in Braun and Kimball, 2021). Palaeognathae phylogeny has received substantial attention (e.g., Harshman et al., 2008; Haddrath and Baker, 2012; Smith et al., 2013; Grealy et al., 2017; Yonezawa et al., 2017; Wang et al., 2022b; Widrig and Field, 2022; Takezaki, 2023) and it has been suggested that this part of the paleognath tree is in the anomaly zone (Cloutier et al., 2019; Simmons et al., 2022). The results of our analyses using the allfam datasets analysis are consistent with the hypothesis that Novaeratitae is an anomaly zone problem and they suggest that MSC methods—including METAL—can exhibit better behavior than ML concatenation under some circumstances.

Several other avian clades have exhibited a dependence on analytical methods and one explanation for their variable recovery in previous analyses would be the hypothesis that they are in (or near) the anomaly zone. The positions of Accipitriformes, Strigiformes, and Coliiformes at the base of Telluraves have varied in earlier studies (Wang et al., 2011; Suh, 2016; Braun et al., 2019; Kuhl et al., 2021; Wang et al., 2022a); some of these conflicts have been suggested to reflect analytical methods (compare the topologies in parts c vs d of Fig. 1). The broader set of methods that we used in this study did not show a clear relationship between analytical method and the topology at the base of Telluraves. Almost all analyses of both allfam datasets supported Accipitriformes+Strigiformes (Figs. 3a and 5a) but only one analysis (wASTRAL) supported Coraciimorphae+Australaves. Recovery of Accipitriformes+Strigiformes in analyses of the Jarvis datasets was also variable (Figs. 7 and 8a), as are the positions of Accipitriformes and Strigiformes within Telluraves (Fig. 9 and Supplementary Fig. S2). However, we note that the recovery of Accipitriformes+Strigiformes sister to Australaves by METAL (node *γ* in Fig. 9a) is congruent with the topology of primary tree in Wu et al. (2024), suggesting that the METAL topology is plausible. On the other hand, the observation that relationships among Accipitriformes, Strigiformes, and other Telluraves vary analyses of the Jarvis datasets (both in this study and in earlier studies, including Jarvis et al., 2014) while being somewhat more consistent in datasets with more extensive taxon sampling (e.g., the allfam dataset) can explained by postulating taxon sampling has a major impact on topology. Regardless, the use of ML concatenation vs MSC methods does not appear to be important for establishing the topology in this part of the tree, falsifying the hypothesis shown in Fig. 1c,d.

Another relationship that has been suggested to differ in a consistent way ML concatenation and MSC trees is the position of Calyptophilidae within the passerine superfamily Emberizoidea (Oliveros et al., 2019b). However, the results of this study were not consistent with the idea that the position of Calyptophilidae differs between ML concatenation trees and MSC trees in a consistent manner. ML concatenation trees identified by different programs (ExaML vs IQ-TREE) placed Calyptophilidae in different positions, one of which was consistent with MSC trees (including the METAL trees).

Both the analyses of the allfam dataset and the reanalyses of the Jarvis datasets provide additional insights into avian phylogeny. The ML concatenation trees for the allfam datasets recover more reliable clades than any other method tested, so clades in the allfam ML concatenation trees that agree with other phylogenomic studies should be viewed as further corroborated. For example, the position of the Neoaves root in the allfam ML concatenation trees is especially provocative; it matches the position of the Neoaves root in Kuhl et al. (2021) and in several trees in Braun and Kimball (2021). This root position is consistent with the METAL tree for the Jarvis intron data (node *δ* in Fig. 9a) and it conflicts with the position in the ML concatenation tree for the same dataset (node *δ* in Fig. 9b). However, analyses of the Jarvis intron data using the gene tree summary methods also revealed the same two conflicting root positions (Supplementary Fig. S2) and the recent phylogenomic analysis by Wu et al. (2024) supported a very different root position. Ultimately, these results might indicate that those nodes with variable recovery in avian phylogenomic trees are the nodes where “noise” generated by to ancient population processes dominate the results of analyses. The apparent good performance of ML concatenation, judged by successful reliable clade recovery, is evidence that the reliable clades—even the hard ones—lie in the part of parameter space where ML concatenation is unbiased. Of course, any attempt to identify reliable clades will by its nature favor clades that are easy. What is interesting is that the intensive study of avian phylogeny has not revealed much, if any, evidence for clades that lie in parts of parameter space where ML concatenation is biased due to the MSC.

The nature of the studies used to define reliable clades is likely to be responsible for our failure to identify reliable clades that fall in the parts of parameter space where ML concatenation is biased (with the possible exception of Novaeratitae). Sayyari and Mirarab (2016) found that “…a branch that appears in only 40% of gene trees can still be resolved with high confidence if a sufficient number of genes are available (e.g., *∼*500).” Most of the phylogenomic studies in Table 1 exceed that number, but the multigene studies do not and they were considered when we defined reliable clades. The number of loci in both allfam UCE datasets should be sufficient based on Sayyari and Mirarab (2016), but the mean alignment lengths for individual loci in those datasets are relatively short (median length of 470 bp in the complete dataset) compared to those in the Jarvis UCE or intron datasets (which have median alignment lengths of 2,482 bp and 4,867 bp, respectively). Thus, the estimated gene trees for the allfam UCE datasets are likely to be of lower quality than those for the Jarvis datasets. Another alternative might be the idea that at least some of the relationships with variable recovery might be associated with gene trees that do not fit the gene tree spectrum expected given the MSC alone. Gatesy and Springer (2022) reanalyzed the Suh et al. (2015) TE insertion dataset and they suggest that the observed conflicts at the base of Telluraves reflect gene flow. Subdivision within ancestral populations also have the potential to cause the gene tree spectrum to deviate from expectation given the MSC (Slatkin and Pollack, 2008). Other relationships in large-scale estimates of avian phylogeny that exhibit variable recovery may reflect the existence of other relationships in the bird tree that do not fit purely treelike evolution; the open question is how much we can infer about the causes of those deviations from expectation when those divergences are relatively ancient, like those at the base of Neoaves where the radiation occurred *>*50 million years ago (based on relaxed molecular clock analyses; e.g., Jarvis et al., 2014; Prum et al., 2015; Kimball et al., 2019).

### MSC analyses and the recombination landscape of genomes

MSC methods have been criticized based on their potential sensitivity to the recombination landscape of genomes (Springer and Gatesy, 2018). Consistency for gene tree summary methods only holds if one uses true gene trees as input but most studies (including this study) use trees estimated from genomic regions without determining whether all sites in a region have the same evolutionary history (i.e., without testing whether the chosen regions correspond to “*c*-genes,” which are non-recombinant regions with a single coalescent history; see Doyle, 1995, 1997). Indeed, when Felsenstein (1981) introduced ML estimation of phylogeny using nucleotide sequences he stated that sites in a region are only expected to be drawn from the same distribution when they are close enough that the recombination rate between those sites is less than 1/2*N*_e_ (where *N*_e_ is the effective population size and the focal taxa are diploid). Simulations have shown that phylogenetic analyses of regions with mixed histories due to intragenic recombination can lead to errors in tree estimation (Schierup and Hein, 2000), and this might be problematic for gene tree summary analyses of the Jarvis datasets, especially the intron data that includes many alignments that correspond to long genomic regions. It is unclear whether this is a problem in typical empirical studies, but Zhao et al. (2023) found that intralocus recombination was a risk factor for low gene tree accuracy. Computationally-efficient phylogenetic methods that are both consistent estimators of the species tree given ILS and insensitive to the existence of short *c*-genes are desirable. Both SVDquartets and METAL have the potential to be such methods, but both performed poorly in this study.

Springer and Gatesy (2018) argued that SVDquartets (and similar methods—which would include METAL) should “…ideally use variation at widely spaced, independently segregating, individual nucleotide positions” and their reasoning is sound when the method is applied to empirical datasets (which are by nature finite). One hypothesis to explain the poor behavior of SVDquartets in this study would be the use of concatenated data rather than unlinked nucleotides. However, we believe that depth in the tree is likely to provide a simpler explanation for its poor behavior. SVDquartets exhibited relatively good recovery of reliable families and superfamilies but poor recovery of orders and superorders (Figs. 3 and 5). Overall, our results were consistent with the hypothesis that SVDquartets exhibits poor performance deep in trees, although it may also be somewhat sensitive to long branch attraction (see Fig. 6c). Our conclusions regarding the poor performance of SVD quartets warrants a note of caution: contracting branches with low support eliminated most conflicts with the ML concatenation tree (Table 4). However, SVDquartets has been shown to yield high support for incorrect relationships under at least some conditions (Shi and Yang, 2017); our results add to the need for caution when interpreting SVDquartets trees.

The expectation that METAL will exhibit good behavior might seem to contradict Schierup and Hein (2000), who found that both NJ and ML yielded trees with topological errors when applied to recombinant regions. However, the proofs of consistency for METAL (Dasarathy et al., 2015; Allman et al., 2019) do not contradict the Schierup and Hein (2000) results; the former applies to data comprising very large numbers of *c*-genes whereas the latter focused on 1,000 bp regions with intralocus recombination (i.e., a case with a limited number of *c*-genes). Distances based on large numbers of unlinked loci should converge on an average that reflects the number of substitutions expected to occur after speciation plus some additional divergence that reflects the variation in the ancestral population (see Edwards and Beerli, 2000, for additional details). Given a variety of reasonable circumstances, this results in distances that will yield the true tree (at least in expectation). The fundamental assumption we have made in this study is that our phylogenomic datasets are large enough for the distance estimates to exhibit good behavior.

### Distance estimators are an important part of METAL analyses

METAL was sensitive to the distance estimators used. This is not surprising, as NJ is only guaranteed to yield the true tree when it is given the true distances as input (Susko, 2004; McTavish et al., 2015). This explains why *p*-distances did not yield accurate estimates of phylogeny even though the low variance of *p*-distances has been suggested to be advantageous (Yoshida and Nei, 2016). Thus, the results conform to our expectation and confirm the importance of using corrected distances (see Braun, 2023, for additional discussion of this issue). Logdet distances typically exhibited the best behavior in this study. The logdet distance estimator is related to the general Markov model (Steel, 1994), so it should be robust to changes in the model of sequence evolution over time (Holland et al., 2012). This is likely to be important because shifts in the model of evolution have been documented in many taxa (including birds; see Berv et al., 2022). However, it has long been known that logdet distances are not robust when there is siteto-site rate heterogeneity (Lake, 1994; Lockhart et al., 1994). In fact, Allman et al. (2019) highlighted a case where NJ of logdet distances is expected to yield an incorrect tree; that example was a classic four-taxon “Felsenstein zone” model tree (i.e., a tree with two separated long terminal branches, a short internal branch, and two short terminal branches) combined with rate heterogeneity. Failure to recover the true tree in that example is unrelated to the MSC, instead it reflects the mixture of site rates and the model tree topology. Removing invariant sites before calculating logdet distances (i.e., logdet-inv distances) has been suggested to address the issue of among-sites rate heterogeneity (Waddell, 1995; Lockhart et al., 1996), but we did not find that this was the case. Instead, the best behavior emerged when we limited the logdet distance calculation to parsimony informative sites, a correction used in Lockhart et al. (1994) but seldom used since. However, even this more radical correction for among-sites rate heterogeneity appeared to yield trees that were affected by long-branch attraction when it was used (e.g., Fig. 6a). This suggests that better ways to address the issue of among-sites rate heterogeneity will be necessary if logdet distances are to become a useful tool in phylogenomics.

Although NJ of logdet inf site distances exhibited the best behavior overall, NJ of trace and ML distances also exhibited good behavior with the Jarvis intron data. The simplest explanation for this difference would be the idea that introns may exhibit fewer (or less extreme) shifts in their model of evolution than UCEs, resulting in better performance for trace and ML distances (which assume a time-reversible model of sequence evolution). This explanation is unlikely to be correct; the Jarvis intron dataset exhibits greater variation in base composition than the Jarvis UCE dataset. Comparisons of base frequencies for different taxa revealed that most of the differences in base composition reflect changes in GC content, which ranges from 39.3%–42.3% (difference = 3%) for the intronic data and 38.1%–39.8% (difference = 1.7%) for the UCE data (see Table S5 for additional information about base frequencies). The differences among taxa in their GC content are relatively modest for both data types, so it seems unlikely that shifts in base composition can explain the different behaviors of logdet vs trace/ML distances. However, even if the observed shifts in base composition are sufficient to be misleading they would be expected to lead to worse behavior for trace and ML distances, since logdet distances should be more robust to model shifts.

We expected introns to exhibit less among-sites rate heterogeneity than UCEs and the ML estimates of the rate heterogeneity parameters confirmed that expectation (*α* = 2.102 without invariant sites for the Jarvis introns vs *α* = 1.17 with proportion of invariant sites = 0.177 for the Jarvis UCEs). The greater among-sites rate heterogeneity that the Jarvis UCE dataset exhibits has the potential to cause logdet distances to behave poorly. We addressed the issue of among-sites rate heterogeneity by limiting the logdet distance calculations to parsimony informative sites distances, but that relatively simple approach is less principled than the use of Γ-distributed rates across sites (which is straightforward with trace and ML distances). Thus, one would predict that trace and ML distances would exhibit good behavior with both data types and logdet distances would only be predicted to perform well only when used with intron data. We observed the opposite of that (see Figs. 7 and 8). Although the basis for the apparent broad utility of logdet inf sites distances is unclear, the apparent utility of such a simple distance estimator in analyses of both UCE and intron data can be viewed as promising. Nevertheless, it seems reasonable to suggest that efforts to develop better distance estimators and find better ways to choose the best distance corrections is important if METAL is to become a useful phylogenomic method.

### Potential to overcome the weaknesses of METAL

We found empirical evidence that METAL has a number of weaknesses, including its sensitivity to missing data and its requirement for large amounts of data. The generality of the missing data problem is unclear; METAL exhibited better performance with both Jarvis datasets than with the complete allfam dataset despite the larger amounts of missing data in the Jarvis datasets (Table 2). This suggests that the problem is the distribution of missing data and further suggests that there are technical solutions for this problem that can be implemented in the laboratory (e.g., expanded use of whole genome sequencing or the use of strategies like tiling in sequence capture to improve data recovery). However, the large amount of data that METAL appears to require represents a bigger problem. Although it is now relatively straightforward to produce data matrices with a size comparable to the Jarvis intron dataset, generating datasets of that size is impossible for organisms with very small genomes (or organisms with insufficient sequence data that can be aligned successfully). Reducing the amount of data necessary for successful METAL analyses, but distance methods intrinsically lose information when aligned sequences are converted into pairwise distances (this has long been recognized, see Penny, 1982). A potential solution to this problem is to consider the correlations among the distances that reflect their shared evolutionary history (Allman and Rhodes, 2010). The idea of using covariances in distance analyses has a very long history (CavalliSforza and Edwards, 1967; Chakraborty, 1977; Felsenstein, 1984; Bulmer, 1991; Roch, 2010), but there have been limited efforts to develop practical implementation of distance-based phylogenetic methods that use covariance information.

The only tree-building method that considers covariances with a practical implementation is BioNJ (Gascuel, 1997), but it uses approximate covariances. We tested BioNJ and found that it generated trees identical to the NJ trees for the same distance estimator (File S2 includes a BioNJ tree). Bulmer (1991) described the use of generalized least squares for phylogenetics, which does consider covariances, but he examined all possible tree topologies making the number of topologies examined too large to be practical for taxon-rich trees. Bulmer (1991) also pointed out that one could test whether distance estimators are adequate, which is more principled than testing a set of distance estimators to find the one that maximizes the number of reliable clades recovered. After all, the reliable clade recovery criterion does not rest on a firm theoretical foundation and it can only be used with phylogenies that have received sufficient study to permit the identification of reliable clades. Generalized least squares can be problematic if the variance-covariance matrix of the distances is singular, which occurs when the distances are small (Rzhetsky and Nei, 1992), but that is unlikely to be a major problem for typical empirical studies. If other distance-based tree building methods are used for METAL analyses it would be desirable to specify the tree building method as well as the distance estimator. However, at this time, it is unclear that there is a practical distance-based tree building method for METAL that is clearly superior to NJ; thus, we believe it is reasonable to focus on NJ as a tree-building method at this time.

## Conclusions

ML concatenation was clearly the best analytical method in comparisons of methods when we used the allfam UCE datasets. We only found one case where MSC methods appeared to correct a bias in ML concatenation (Novaeratitate) and all MSC methods appeared to be problematic based on the reliable clade recovery criterion. SVDquartets appeared to behave especially poorly deep in tree. A simple explanation for the observation that ML concatenation generally performed better than other methods would be to postulate that discordance among gene trees is not primary source of topological errors for the clades we could assess. Other issues, like robustness to long branch attraction, might be more important issues. All methods exhibited better behavior when they were used to reanalyze the Jarvis datasets, but even here there was no indication that ML concatenation was problematic. Unlike the case for the allfam datasets, where we had limited *a priori* information regarding the potential for any analysis to resolve many relationships using those data, we expected ML concatenation to exhibit relatively good behavior with the Jarvis datasets. After all, the published ML concatenation topologies were known to include most reliable clades and it was very unlikely that our reanalyses using ML concatenation would yield trees that differed from those in Jarvis et al. (2014) in fundamental ways. The Jarvis datasets were also used in the papers describing wASTRAL and wASTRID (Zhang and Mirarab, 2022; Liu and Warnow, 2023), so there was already evidence that those methods can recover many reliable clades (although neither of those studies analyzed UCEs or introns separately). Thus, the primary questions we could ask using the Jarvis datasets was whether SVDquartets and/or METAL would exhibit good performance. The larger numbers of sites did not improve the performance of SVDquartets, but it did improve the performance of METAL. However, our empirical results suggested that METAL is less promising than one might believe based on theory (Dasarathy et al., 2015; Allman et al., 2019) or simulations (Rusinko and McPartlon, 2017; Wilson and Rogers, 2023). Despite the disappointing results, it seems reasonable to suggest that METAL warrants additional research; after all, it is not computationally demanding and there are reasons to believe that its performance can be improved. Looking beyond the information about method performance, our new family-level estimates of avian phylogeny are likely to be useful to the community. As long as one views the poorly supported parts of the tree with appropriate caution there is every reason to view the ML concatenation tree for the complete allfam dataset (Fig. 2) as an accurate estimate of avian phylogeny (albeit one with a small number of biased branch length estimates).

## Appendix 1 Distance estimators

Although the first distance estimator was described more than 50 years ago (Jukes and Cantor, 1969), most of the development of distance estimators for phylogenetics occurred in the 1980s. Many distance estimators have a clear link with specific models of sequence evolution that are used in ML analyses, but that link is less clear in other publications that describe distance estimators. For example, Tajima and Nei (1984) described the formula for the F81 model of sequence evolution (Felsenstein, 1981), but they did not directly link their distance to the model for ML estimation and they also presented three different treatments of the base frequency parameters. The formula that Tajima and Nei (1984) favored treats base frequencies differently from the simplest treatment for the F81 model. This history makes the best nomenclature for some distance estimators unclear. We use the term trace distances for the general distance formulae that have been derived repeatedly (e.g., Lanave et al., 1984; Rodríguez et al., 1990) in different forms; the name reflects the Rodríguez et al. (1990) formula shown in equation 1:

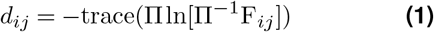

where *d*_*ij*_ is the genetic distance for sequences *i* and *j*, Π is a diagonal matrix of nucleotide frequencies, and F_*ij*_ is the divergence matrix for the sequence pair. The Lanave et al. (1984) formula is different, but Waddell (1995) showed that it yields distance estimates that are identical to equation 1, suggesting algebraic identity. Zharkikh (1994) also showed that the Lanave et al. (1984) formula provides distances for the GTR model, further emphasizing the relationship between the Rodríguez et al. (1990) formula and the GTR model. Swofford et al. (1996) noted that it was possible to modify equation 1 to produce distance estimators for any submodel of the GTR model. Waddell (1995) and Waddell and Steel (1997) described a way to incorporate among-sites rate heterogeneity into the distance estimator, by substituting the inverse of the moment generating function for the Γ distribution for the matrix logarithm, yielding equation 2:

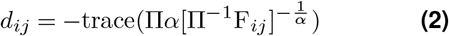

where *α* is the Γ distribution shape parameter (other distributions could be used to describe among sites rate heterogeneity; for details see Waddell and Steel, 1997). The Π and F_*ij*_ matrices can be calculated from pairwise comparisons, but the *α* parameter requires more than two sequences to be identifiable (Steel, 2009). We used ML concatenation to estimate *α* in this study.

The nomenclature for the other distances estimators that we used is much clearer. *p*-distances are simply normalized Hamming distances. ML distances are calculated from pairs of aligned sequences by optimizing the “branch length” for a two taxon tree given some model of sequence evolution. ML distances are expected to yield values close to those for trace distances when they are calculated using the same model; the difference is that the rate matrix is fixed at user specified values for ML distances and estimated from the pair of sequences for trace distances. ML distances are likely to be superior to trace distances for short sequences, where one can reduce the variance of the rate matrix parameters by generating them by ML concatenation or some other method (e.g., Tamura et al., 2004). On the other hand it seems reasonable to hypothesize that trace distances provide better distance estimates when sequences are long because it may be able to capture variation in the model across the tree (although we note that the GTR model has properties that may be problematic when there are model shifts; see Sumner et al., 2012). The relationship between the logdet distance estimator and the general Markov model (Steel, 1994) suggests that it will be less problematic when the model of sequence evolution changes over the tree.

## Appendix 2 Bird images

The bird photographs in Fig. 2 were taken from Wikimedia Commons. The images were available under the Creative Commons Attribution-Share Alike 2.0 Generic license (images 5–7 and 9) or the Creative Commons Attribution-Share Alike 4.0 International license (images 1–4 and 8). The only modification to the images was cropping to make the birds more visible relative to the background. Links to the original images and the photographers that generously made the images available are: 1) *Picathartes gymnocephalus*, Charles J. Sharp (image, photographer); 2) *Sarcoramphus papa*, Chris Down ((image, photographer); 3) *Alca torda*, Chme82 ((image, photographer); 4) *Topaza pella*, author: Kyle Van Houtan (image, photographer); 5) *Aramus guarauna*, Gary Leavens (image, photographer); 6) *Crinifer piscator*, author: peterichman (image, photographer); 7) *Otidiphaps nobilis*, Mark Dumont (image, photographer); 8) *Pelecanus occidentalis*, Frank Schulenburg (image, photographer); and 9) *Podiceps auritus*, Ekaterina Chernetsova (Papchinskaya) ((image, photographer).

## Acknowledgements

This work was supported by the United States National Science Foundation (grants DEB-1655683 to E.L.B. and R.T.K. and DEB-1655624 to B.C.F. and R.T.B), the Smithsonian Institution’s Scholarly Studies Program (to M.J.B, E.L.B., T.C.G., R.T.B., and B.C.F.), and the Smithsonian’s Grand Challenges Program (Consortium for Understanding and Sustaining a Biodiverse Planet) (to M.J.B.). We thank the following institutions and individuals for loaning tissues and DNA samples: American Museum of Natural History (AMNH), Academy of Natural Sciences, Philadelphia (ANSP, Australian National Wildlife Collection (ANWC), Field Museum of Natural History (FMNH), University of Kansas, Museum of Natural History (KU), Llewellyn D. “Lou” Densmore, Texas Tech University (LLD-TTU), Louisiana State University Health Sciences Center, New Orleans, courtesy of Herbert C. Dessauer (LSU-HSC), Louisiana State University, Museum of Natural Science (LSUMZ, Museums Victoria, Australia (MVIC), Museum of Vertebrate Zoology, Berkeley (MVZ), San Francisco Zoo & Gardens (SFZG), Smithsonian National Museum of Natural History (USNM, Burke Museum of Natural History and Culture, University of Washington (UWBM), and Universitets København, Zoologisk Museum (ZMUC).

## Supporting Information

Supplementary Materials available from the Zenodo Digital Repository: https://doi.org/10.5281/zenodo.10836298 and from GitHub: https://github.com/ebraun68/METALtest

## Notes

https://github.com/ebraun68/METALtest

https://doi.org/10.5281/zenodo.10836298

### Competing Interest Statement

The authors have declared no competing interest.

### Summary of Updates

This manuscript includes an assessment of "reliable clades" in avian phylogeny. This revision better incorporates the findings of another large-scale phylogenomic tree of birds (Wu et al. https://doi.org/10.1073/pnas.2319696121). The new data are incorporated as Figure S1.

